# Ran-GTP assembles a specialized spindle structure for accurate chromosome segregation in medaka early embryos

**DOI:** 10.1101/2023.07.20.549960

**Authors:** Ai Kiyomitsu, Toshiya Nishimura, Shiang Jyi Hwang, Satoshi Ansai, Masato T. Kanemaki, Minoru Tanaka, Tomomi Kiyomitsu

**Affiliations:** Okinawa Institute of Science and Technology Graduate University, 1919-1 Tancha, Onna-son, Kunigami-gun, Okinawa 904-0495, Japan; Division of Biological Science, Graduate School of Science, Nagoya University, Chikusa-ku, Nagoya 464-8602, Japan; Hokkaido University Fisheries Sciences, 3-1-1, Minato-cho, Hakodate, Hokkaido 041-8611, Japan; Graduate School of Life Sciences, Tohoku University, Sendai, Miyagi 980-8577, Japan; Laboratory of Genome Editing Breeding, Graduate School of Agriculture, Kyoto University, Sakyo-ku, Kyoto 606-8507, Japan; Department of Chromosome Science, National Institute of Genetics, Research Organization of Information and Systems (ROIS), and Graduate Institute for Advanced Studies, SOKENDAI, Yata 1111, Mishima, Shizuoka 411-8540, Japan; Department of Biological Science, The University of Tokyo, Tokyo 113-0033, Japan

## Abstract

Despite drastic cellular changes during cleavage divisions, a mitotic spindle is assembled in each blastomere to accurately segregate duplicated chromosomes. Recent studies indicate that early embryonic divisions are highly error-prone in bovines and humans. However, processes and mechanisms of embryonic spindle assembly remain little understood in vertebrates. Here, we established live functional assay systems in medaka fish (Oryzias latipes) embryos by combining CRISPR knock-in with an auxin-inducible degron technology. In contrast to mammals, mitoses during cleavage divisions are very rapid (<12 min), but segregation errors are rarely observed. Importantly, we found that the Ran-GTP gradient assembles a specialized, dense microtubule network at the spindle midplane during metaphase, which is essential for faithful chromosome segregation in early embryos. In contrast, Ran-GTP becomes dispensable for chromosome segregation in later stages, where spindles are morphologically remodeled into short, somatic-like spindles lacking the dense microtubule network. We propose that the specialized Ran-based spindle structure ensures high fidelity of chromosome segregation in large, vertebrate early embryos.

## Introduction

During early embryogenesis in animals, a large, fertilized egg undergoes repeated cell divisions called cleavage divisions to create numerous small, differentiated cells^1^. This process comprises a series of dynamic physical and biochemical changes, including cell size reduction^2–7^, zygotic gene activation^8–11^, and cell cycle remodeling^12, 13^. Regardless of these drastic cellular changes, unified parental chromosomes must be accurately duplicated and segregated to all blastomeres to maintain and transmit genomic information. Although sizes and cleavage patterns of fertilized eggs vary among species^14^, a microtubule-based bipolar structure, the mitotic spindle^15^, is generally assembled in each blastomere to segregate duplicated chromosomes into daughter cells. Recent studies have shown that embryonic divisions in bovine and human are error-prone^16, 17^, but mechanisms of spindle assembly and chromosome segregation in vertebrate embryos remain poorly understood, compared to those in somatic cells.

In the 1980s, based on the dynamic properties of microtubules (MTs), Kirschner and Mitchison proposed a simplified search and capture model^18, 19^ for mitotic spindle assembly. In this model, centrosomes act as a major MT organizing center, and dynamic MT plus-ends are captured by kinetochores on chromosomes to form a bipolar spindle. On the other hand, in the 1990s, Heald and colleagues proposed a self-organization model using *Xenopus* egg extracts^20, 21^. In this model, chromatin acts as a MT nucleation site, and MTs around chromosomes are subsequently coalesced and focused into spindle poles in the absence of centrosomes. Afterward, several studies established the key concept that conserved chromatin-bound RCC1 (regulator of chromosome condensation 1) generates a spatial Ran-GTP gradient^22–27^, which promotes self-organization of the spindle by locally activating spindle assembly factors (SAFs) near chromosomes^28–30^. Consistent with this model, the Ran-GTP gradient is essential for acentrosomal spindle assembly in female meiosis^31–33^. However, we recently found that whereas this gradient affects localization of some of SAFs, the Ran-GTP gradient is dispensable for bipolar spindle assembly in centrosome-containing somatic human cells^34^, as observed in other somatic cell lines^35, 36^, probably due to functions of centrosomes and other pathways.

What function does Ran-GTP serve in spindle assembly in large, centrosome-containing vertebrate embryonic cells? Considering the large distance between chromosomes and centrosomes (> 20 μm in *Xenopus laevis* early embryos^4, 37^), Ran-GTP may have unique essential functions for embryonic spindle assembly, despite the presence of centrosomes. Interestingly, in *Xenopus laevis* embryonic extracts, Ran-GTP is required for spindle assembly at the 4-cell stage, but not at the ∼4,000 cell stage^3^. However, it remains unclear how Ran-GTP promotes spindle assembly and chromosome segregation in live *Xenopus* embryos and in other species. In addition, even though mechanisms of spindle scaling, including the upper limit to mitotic spindle length have been well investigated^3, 4, 6, 7, 38^, it remains unclear whether and how early embryos assemble a specialized spindle structure for accurate chromosome segregation during cleavage divisions.

Compared to mammals and frogs, fish embryos are suitable for live imaging. Since these transparent embryos show meroblastic cleavage^14^, spindle dynamics can be relatively easily analyzed^7, 39–41^ throughout embryogenesis. Compared to zebrafish, Japanese medaka, *Oryzias latipes*, has several advantages, including smaller genome size^42^, a wide range of permissive temperatures, and daily egg production^43^, which are helpful for genome editing and handling of embryos. In this study, we analyzed dynamic mechanisms of spindle assembly and chromosome segregation in medaka early embryos by combining high-quality live imaging with CRISPR/Cas9-mediated genome editing and an auxin-inducible degron 2 (AID2)-based protein knockdown system^44^. Our live functional studies revealed that Ran-GTP is required for accurate chromosome segregation by assembling a dense MT network around the metaphase spindle midplane, specifically in early embryos.

## Results

### Dual-color live imaging of chromosomes and microtubules in medaka early embryos

Medaka early embryonic development was carefully observed using a stereo microscope^2^. To investigate intracellular dynamics, we first visualized chromosomes and MTs by generating transgenic medaka expressing RCC1-mCherry2 (RCC1-mCh) and EGFP-α-tubulin (Figure 1A and 1B). mCherry2 coding sequence (CDS) was integrated into the C-terminal region of the endogenous RCC1 gene using 5’ modified dsDNA donors^45^ (Figure S1A), whereas CDS of EGFP-α-tubulin was inserted at the 5’ UTR of the tubulin alpha-1B gene on Chromosome 7 (Figure S1B) (see Methods for details). Homozygous adult medaka having both RCC1-mCh and EGFP-α-tubulin grew normally, and fertilized eggs obtained from homozygous pairs were used for live imaging (Figure 1B).

**Figure 1.**
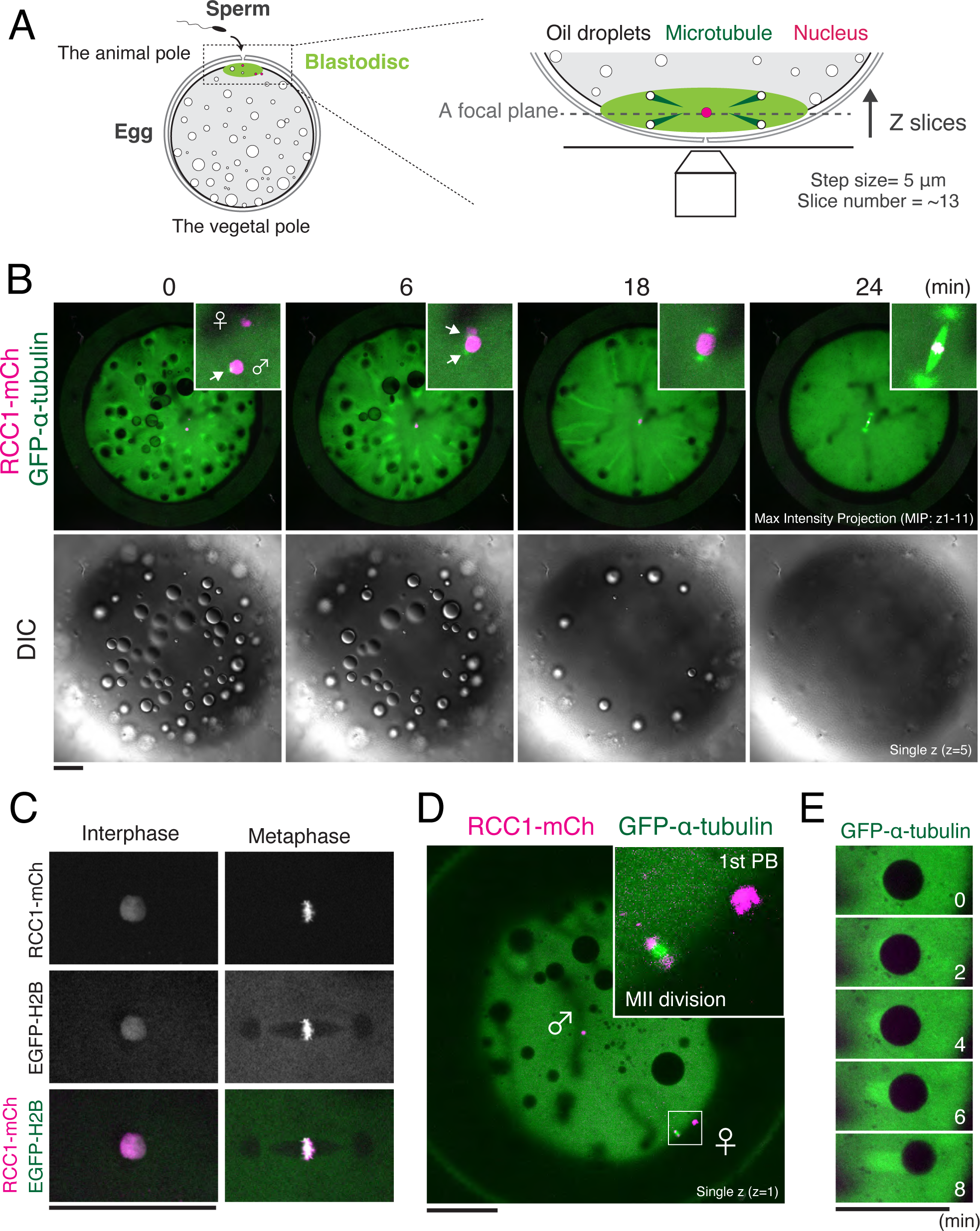
Live imaging reveals dynamic behavior of chromosomes and MTs in medaka fertilized eggs. (A) Schematic representation of microscope observation of a medaka fertilized egg. (B) Live-embryo images showing dynamics of chromosomes (magenta), MTs (green), and oil droplets. Maximal-intensity-projection (MIP) images are shown. Insets indicate pronuclear fusion and a zygotic spindle. White arrows indicate centrosomes. (C) Live-cell fluorescent images showing co-localization of RCC1-mCh with EGFP-Histone H2B. (D) A single-plane live-cell fluorescent image showing the first polar-body (PB) and Meiosis II division at the periphery of the blastodisc. (E) Representative GFP-α-tubulin fluorescent images showing generation of comet-like structure under the oil droplet. Scale bars = 100 μm.

RCC1 is a well-conserved chromatin-binding protein^46–48^ (Figure S1C). RCC1-mCh co-localized with Histone H2B (a marker of chromosomes) (Figure 1C), and visualized Meiosis II chromosomes and polar bodies (PBs) as well (Figure 1D). After fertilization, a female pronucleus migrated (Figure 1B t=0) toward a male pronucleus with centrosomes, locating around the center of the blastodisc near the animal pole (Figure 1A, 1B and 1D). In contrast, oil droplets were displaced from the animal pole to the vegetal pole before the first mitosis^2^ (Figure 1B). This oil droplet migration is well known, but the driving force is unclear. Interestingly, comet-like structures of EGFP-α-tubulin were formed under these oil droplets (Figure 1E) and elongated radially during oil droplet displacement (Figure 1B, t=0-18). These results show that dual-color, live imaging is a powerful method to reveal uncharacterized intracellular dynamics in embryogenesis.

### Characterization of medaka cleavage divisions

Spindle assembly and cell cycle remodeling during early embryogenesis have not been carefully analyzed in medaka. To characterize these processes, we next performed long-term, live-cell imaging of fertilized medaka eggs at 3-min intervals for ∼10 hr at 24-25 °C. Metaphase spindles were visualized from the 1-cell to late blastula stage (Figure 2A, S2A and Supplemental Movie S1). Despite long-term imaging, all embryos developed normally to nearly the hatching stage, suggesting very low phototoxicity of these imaging conditions.

**Figure 2.**
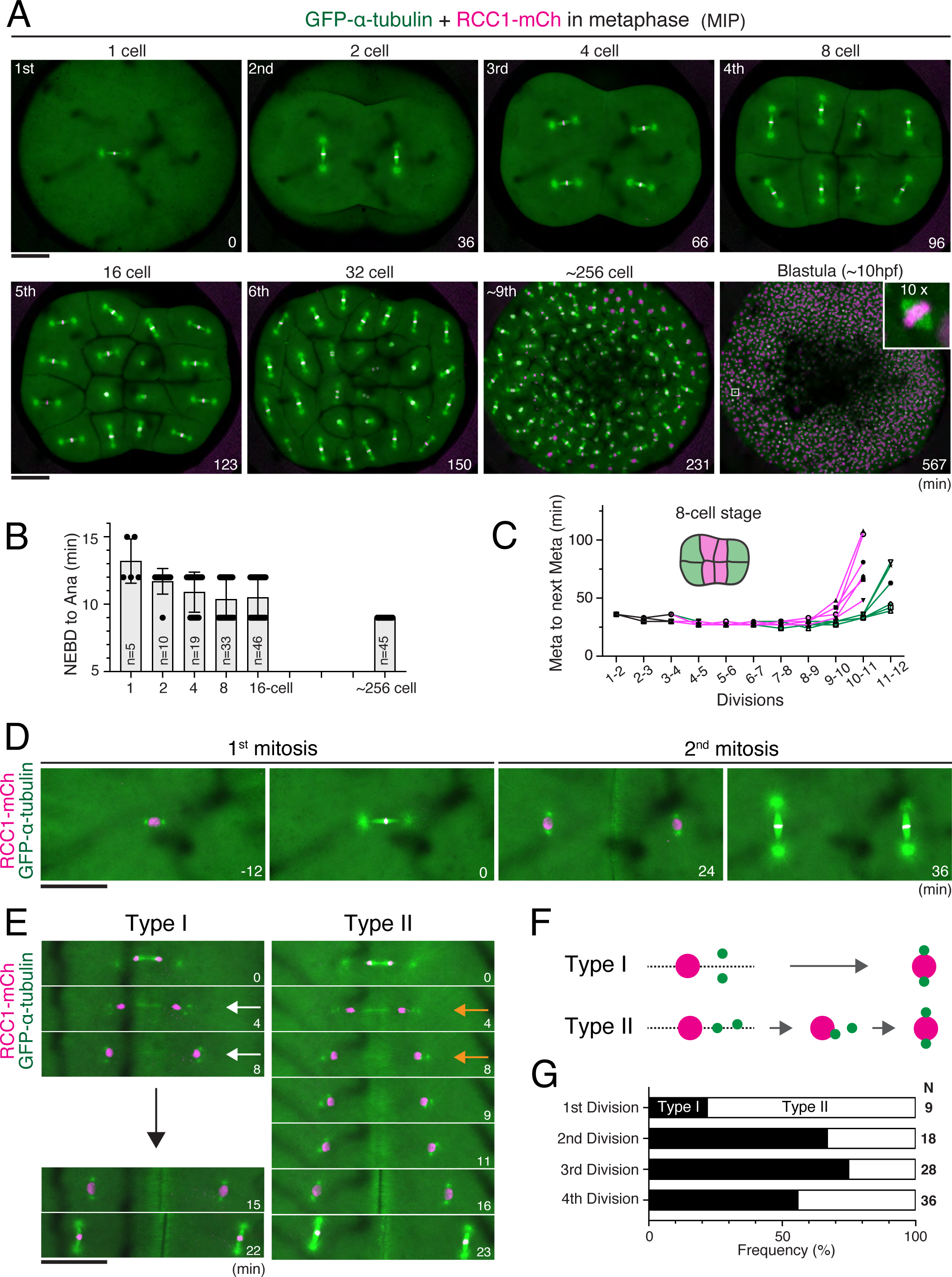
Live imaging shows dynamic regulation of the cell cycle and spindle positioning during medaka cleavage divisions. (A) Representative live-cell images showing metaphase spindles in indicated embryonic stages. Maximal-intensity-projection (MIP) images are shown. (B) Quantification of time elapsed from NEBD to anaphase onset in each stage. Indicated numbers of blastomeres within focal planes were measured in 5 embryos. (C) Quantification of time from one metaphase to the next. Six cell-division lineages were tracked and measured for inner (magenta) and peripheral (green) blastomeres, respectively, from 4 embryos. (D) Live-cell images showing the correlation between centrosome and spindle orientation in the first and second mitosis. (E) Centrosomes are initially separated either vertically (Type I, white arrows) or horizontally (Type II, orange arrows) after mitosis. Centrosomes in Type II cells subsequently oriented vertically before entering the next mitosis. (F) Classification of centrosome separation patterns based on (E). (G) Quantification of centrosome separation types in early embryonic divisions. Error bars indicate mean ± SD. Scale bars = 100 μm.

We first measured mitotic duration from nuclear envelope breakdown (NEBD) to anaphase onset. In contrast to the longer duration in mammalian zygotes (> 60 min in mice^5^, bovines^16^, and humans^17^), the first division in medaka took only ∼12 min and the duration decreased further to ∼9 min at the 256-cell stage (Figure 2B). Despite the short duration, chromosome segregation errors were rarely observed during the first 4 divisions (Movie S1, see also Fig. 4A, 4G and S4B-D for details). The cell cycle length (duration between successive metaphases) was ∼36 min from the 1^st^ to the 2^nd^ division, and the length was constantly ∼30 min until the 9^th^ division (256-cell stage) (Figure 2C), which is similar to the duration in zebrafish (∼15 min) and *Xenopus* (∼30 min), but shorter than that in mice (∼12 hr)^12^. The inner blastomeres of 8-cell stage embryos (Figure 2C) appeared to lengthen the cell cycle at the ∼512-1024 cell stage (10-11^th^ divisions), one round earlier than the peripheral blastomeres (Figure 2C). The timing of this cell-cycle increase seems to be roughly coupled with the timing of zygotic gene activation^8^ (Figure S2A).

Zygotic spindles were oriented horizontally to achieve planar divisions until the 16-cell stage^2^, but spindles in the central four blastomeres or other relatively smaller cells tended to orient perpendicularly during the 5^th^ division (16-cell stage metaphase, Figure 2A, t=123). Anaphase entry was well synchronized until the 16-cell stage, but some smaller cells entered anaphase earlier, at around the 32-cell stage (Figure 2A, t=150). Many cells were not synchronized around the late morula stage (256-512 cell stage, Figure 2A, t=231), after which blastomeres started to migrate in concert with cell cycle elongation (Figure 2C and S2A). Although abnormal divisions were rarely observed until the early morula (64-128 cell) stage, late blastula embryos displayed several polyploid-like nuclei (Figure S2B), which seemed to be generated by cell fusion during the late morula stage (Figure S2C). Together, these results provide fundamental information about cell cycles in medaka embryology.

### Centrosome positioning in medaka early embryonic divisions

Mitotic spindle position and orientation are determined by the position of interphase centrosomes in early frog and fish embryos^39^. Since spindle orientation is well regulated during the first 4 divisions in medaka embryos (Figure 2A), we next analyzed centrosome positioning. Centrosomes, marked by punctate tubulin signals, were always positioned at opposite sides of the nucleus before mitotic entry (Figure 2D t=-12, 24). After chromosome segregation, centrosomes were duplicated in the cytoplasm and separated vertically to the spindle elongation axis (Figure 2E t=4, 2F, Type I), consistent with a previous study^39^. Daughter nuclei subsequently migrated between the separated centrosomes (Figure 2E t=8) before the next mitosis. In some cases, however, centrosomes were transiently separated horizontally (Figure 2E-2G Type II, t=4), but finally became oriented vertically to the spindle elongation axis (Figure 2E and 2F, Type II). The frequency of Type II orientation was relatively high in the first division, but reached ∼50% in the fourth divisions (Figure 2G). These results suggest that centrosome positioning is regulated by multiple mechanisms to ensure proper mitotic spindle orientation in medaka cleavage cycles.

### Unique spindle architecture in medaka early embryos

Since mitotic spindle size scales with cell size in vertebrate early embryos^4, 5, 7, 37^, we next measured medaka metaphase spindles during different stages (Figure 3A-E). Spindle length, defined as centrosome-centrosome distance (solid lines in Figure 3A), decreased significantly from ∼70 to 10 μm (Figure 3C), whereas spindle width and metaphase plate length were relatively constant until the early blastula stage (Figure 3D and S3A). The first spindles were the longest, but thinner compared to the second spindles (Figure 2D and 3D) and were sometimes bent (Figure S3B). Consistent with previous studies^4, 5, 7, 37^, spindle length scaled with cell diameter after the 16-cell stage, but appeared to reach an upper limit in the first to fourth (1-cell to 8-cell) divisions (Figure 3E).

**Figure 3.**
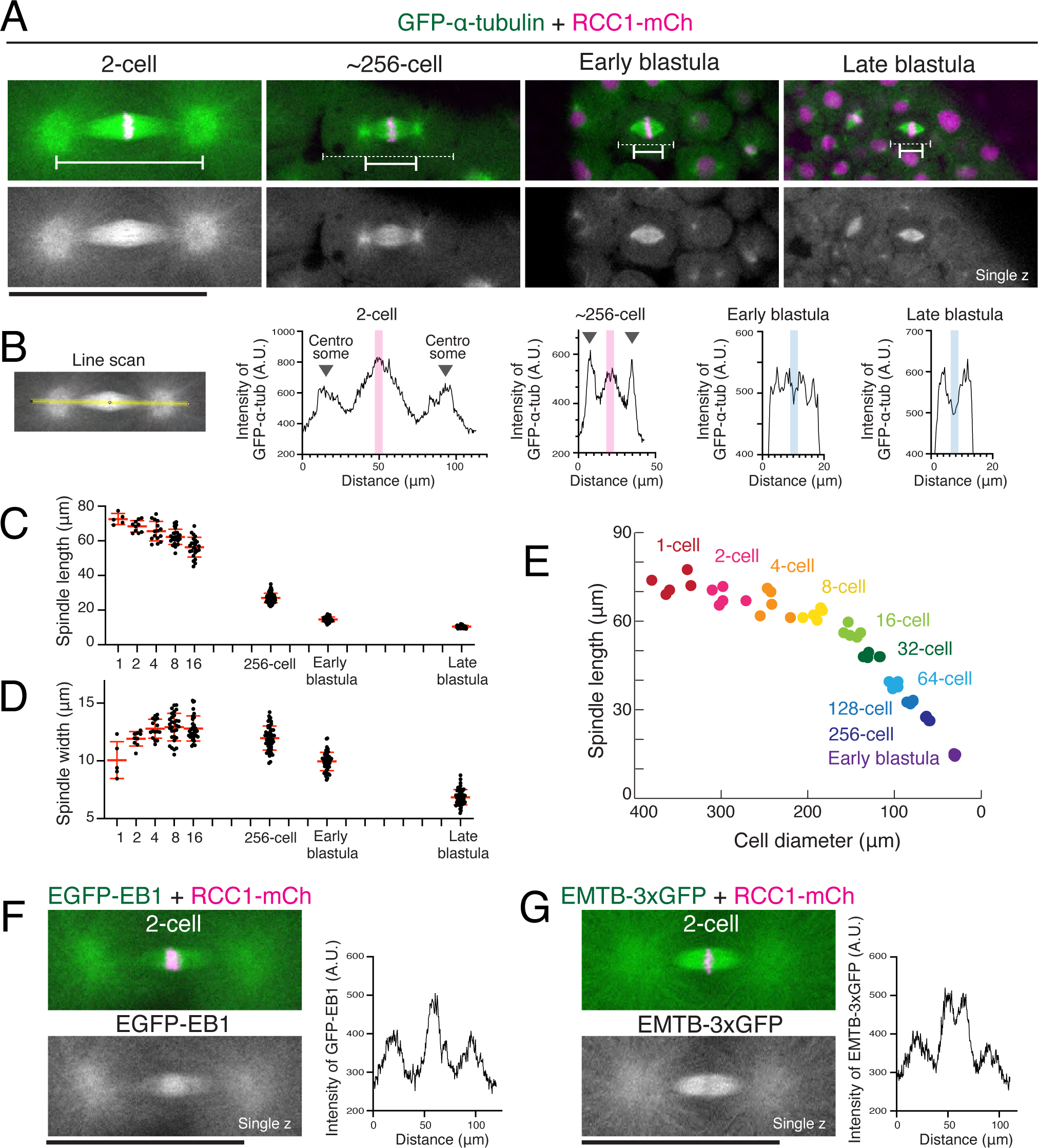
Mitotic spindles display dynamic structural changes during medaka cleavage divisions. (A) Representative live-cell images showing metaphase spindles in indicated embryonic stages. Solid and dashed lines indicate centrosome-centrosome distance and cell diameter, respectively. (B) Left: fluorescence intensity on the yellow line was measured using Fiji. Right: graphs of relative fluorescence intensities for line scans of spindles in (A) showing an increase of EGFP-α-tubulin intensity at the spindle midplane in 2-cell and ∼256-cell, but not in early and late blastula spindles. (C, D) Quantification of spindle length (C) and width (D) in indicated stages. N= 5, >9, >15, >24, >24, >50, >50, >50 for 1-, 2-, 4-, 8-, 16-, 256-, early blastula, and late blastula stages, respectively, from 5 different embryos. (E) Cell diameter and spindle length are plotted as colored circles for individual embryos at different stages, showing spindle length scales with cell diameters after 16-cell stages. These reach an upper limit in the fourth divisions. (F, G) Live metaphase-cell images (left) and a graph for the line scan (right) of GFP-EB1 (F) and EMTB-3×GFP (G), showing accumulation of GFP-EB1 and the relative decrease of EMTB-3×-GFP around the spindle midplane. Error bars indicate mean ± SD. Scale bars = 100 μm.

**Figure 4.**
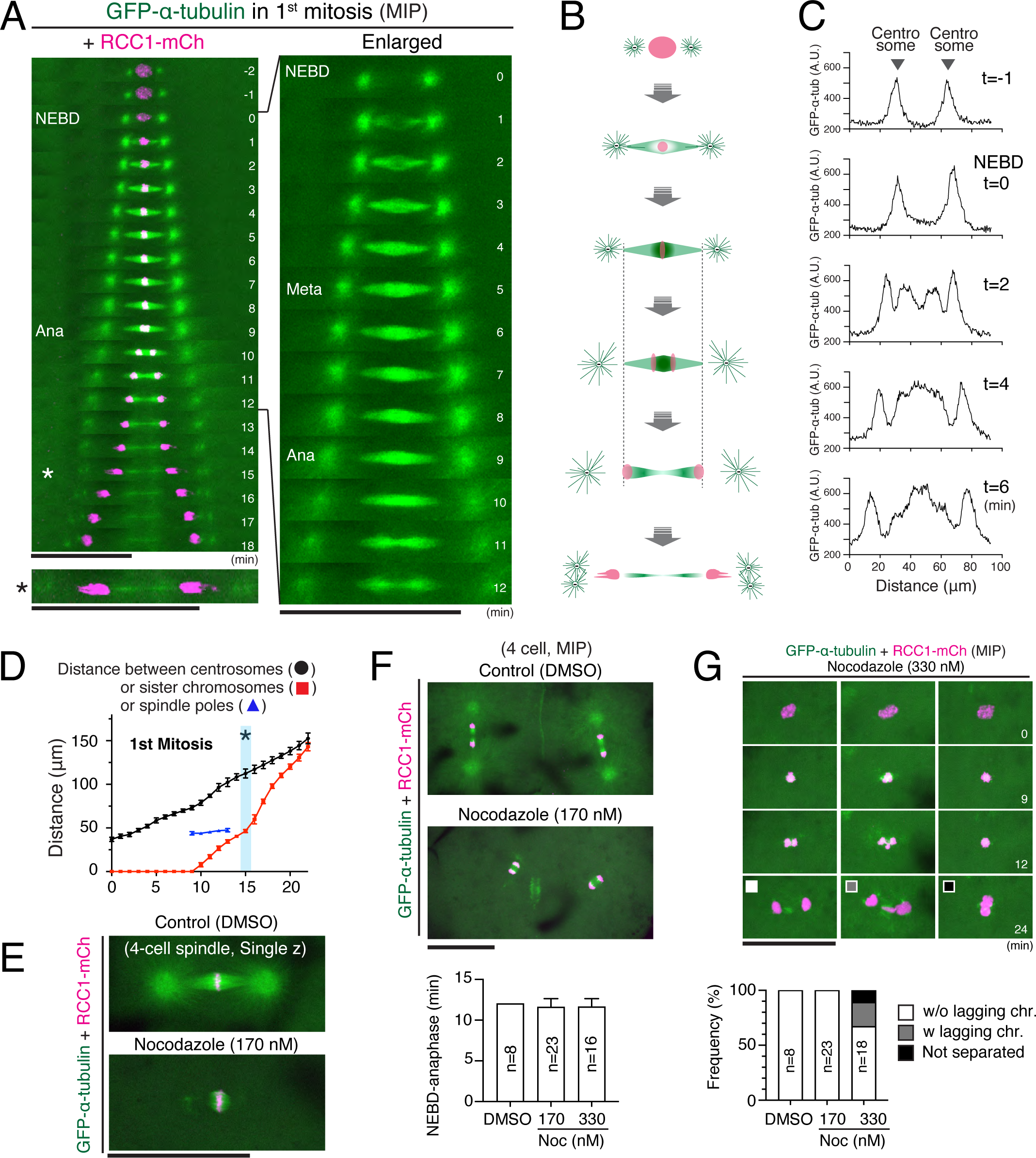
A dense MT network is assembled at the spindle midplane during metaphase in medaka early embryos. (A) Kymographs showing the first mitotic spindle assembly process in a medaka embryo. Asterisks indicate a telophase chromosome or deformed nucleus migrating to centrosomes. (B) Schematic representation of spindle assembly and chromosome segregation processes in (A). Spindle length is almost constant during anaphase. (C) Graphs of relative fluorescence intensities for line scans of spindles in (A). Single z-section images shown in Figure S4A were used for quantification. (D) Quantification of centrosome-centrosome (black), separating chromosomes (red), and pole-pole distance (n=4) during the first mitosis. (E, F) Live-cell images of control (top) and Nocodazole-treated (bottom) 4-cell spindles in metaphase (E) and anaphase (F). The bottom graph in (F) indicates the duration of mitosis in DMSO or Nocodazole-treated 4-cell blastomeres. (G) Top: 330 nM Nocodazole-treated 4-cell blastomeres showing fewer MTs around metaphase chromosomes and abnormal chromosome segregation. Bottom: Quantification of abnormal segregation phenotypes. Error bars indicate mean ± SD. Scale bars = 100 μm.

Strong astral MTs were clearly visible around centrosomes in early embryonic spindles (Figure 3A and 3B). However, their radial signals were gradually reduced and hardly visible after early blastula stages (∼1024-cell stage, Figure 3A, 3B and S3C). Unexpectedly, early embryonic spindles have another MT-dense region around metaphase chromosomes (Figure 3A and 3B, indicated in magenta), which also gradually diminished and became undetectable after early blastula stages (Figure 3B). Exogenously expressed EGFP-EB1, which recognizes growing MT plus-ends^49^, also accumulated around chromosomes in 2-cell spindles (Figure 3F), further confirming that a specialized MT network is formed around chromosomes in early spindles. In contrast, EMTB-3×GFP, which contains the MT-binding domain of ensconsin^50^, localized on spindle MTs more uniformly and showed reduced fluorescence intensities on metaphase plates (Figure 3G). EMTB-3×GFP clearly visualized astral microtubules, but also displayed MT-like signals around cortical cell membranes (Figure S3D), which were rarely observed with EGFP-α-tubulin (Figure 2A), suggesting that this construct may stabilize MTs and/or preferentially recognize stabilized MTs in medaka embryos. Together, these results revealed that medaka early embryonic spindles have unique structures that consist of MTs with different characters and that are remodeled in blastula stages.

### Early embryonic spindles form a dense MT network around chromosomes during metaphase

The above results indicate that medaka early embryonic spindles have two major MT nucleation sites, centrosomes and chromosomes. To understand how these sites promote embryonic spindle assembly, we next performed live-cell imaging at 1-min intervals. After NEBD, a vague spindle-like structure was assembled between the two centrosomes within 1 min and it surrounded chromosomes (Figure 4A t=1-2, 4B and 4C, S4A). GFP-α-tubulin signals were relatively high around the poles, but were still weak around chromosomes (Figure 4A and 4C t=2). However, MTs gradually enriched throughout the spindle during prometaphase (t=3-4) and finally formed a dense MT region around chromosomes during metaphase (t=5-8). Interestingly, this region remained between separated chromosomes after anaphase onset (t=9-10), suggesting that these signals do not simply represent bundled-kinetochore microtubules, but that they also contain bridging or midplane-crossing microtubules^51–53^.

Unexpectedly, we also found that early embryonic spindles have no obvious anaphase spindle elongation, as the pole-pole distance remained almost constant (Figure 4A t=9-12, 4D) like acentrosomal spindles in moss^54^. In contrast, the centrosome-centrosome distance increased gradually after NEBD, finally resulting in a large gap between spindle poles and centrosomes during metaphase (Figure 4A and 4D t=5-8) and anaphase (t=9-12). Once separated chromosomes reached the poles (t=12), decondensed chromosomes or early nuclei migrated toward duplicated centrosomes with transient deformed structures (indicated by asterisks in Figure 4A t=15 and 4D). The same features were observed in the second, third and fourth mitoses (Figure S4B-D). Since centrosomes are spatially separated from the spindle midplane in metaphase, our results indicate that chromosomes contribute significantly to formation of the dense MT network at the spindle midplane.

### MTs around chromosomes are stable and sufficient for chromosome separation

Kinetochore-MTs (k-fibers) are stabilized to drive chromosome alignment and segregation^15^. To understand MT stability of early embryonic spindles, we next treated embryos with 170-nM nocodazole, a MT-destabilizing drug. As expected, the majority of spindle MTs, including astral MTs, were destabilized (Figure 4E). However, MTs around chromosomes were still visible (Figure 4E) and sufficient to segregate chromosomes with normal timing (Figure 4F), although spindle position and orientation were dysregulated at the 4-cell stage (Figure 4F). Treatment with 330-nM nocodazole further destabilized MTs (Figure 4G t=9), which impaired metaphase chromosome alignment and anaphase chromosome segregation (Figure 4G). However, mitotic delay was not observed (Figure 4F), suggesting that medaka early embryos have no functional spindle assembly checkpoint. In summary, these data indicate that MTs around chromosomes are stabilized and sufficient for chromosome segregation, whereas centrosomes and astral MTs are required to control spindle position and orientation in medaka early embryonic divisions.

### Expression of the Ol-RanT27N dominant negative mutant disrupts the dense MT network and causes abnormal chromosome segregation

A chromosome-derived Ran-GTP gradient has been thought to locally activate SAFs by dissociating inhibitory importins from SAFs around chromosomes^28–30, 34^ (Figure 5A). To understand the requirement of Ran-GTP for embryonic spindle assembly in medaka, we next expressed the dominant negative mutant of medaka *Oryzias latipes* Ran, Ol-RanT27N, which corresponds to mouse and human RanT24N (Figure S5A). Exogenous expression of mCherry-tagged wild-type Ol-Ran (mCh-Ol-RanWT) had no obvious effects on the spindle assembly at the 8-cell stage (Figure 5B and S5B). In contrast, expression of the Ran mutant, which binds endogenous RCC1 and inhibits RCC1’s GEF activity^55^, caused severe chromosome segregation defects (Figure 5B and S5B) in an expression-level-dependent manner (Figure S5C). Importantly, Ol-RanT27N expression impaired both spindle assembly timing and spindle architecture. Three min after NEBD, spindle MTs were hardly detectable between centrosomes (Figure 5B and S5C t=3). Subsequently, a faint spindle structure was assembled, but it lacked the dense MT network around chromosomes (Figure 5B, 5D and S5C t=6). Chromosomes indicated by mCh-Ol-RanT27N were abnormally elongated after anaphase and finally pinched or cleaved by a cleavage furrow (Figure 5B and S5C t=15), similar to the fission yeast *cut* phenotype^56^.

**Figure 5.**
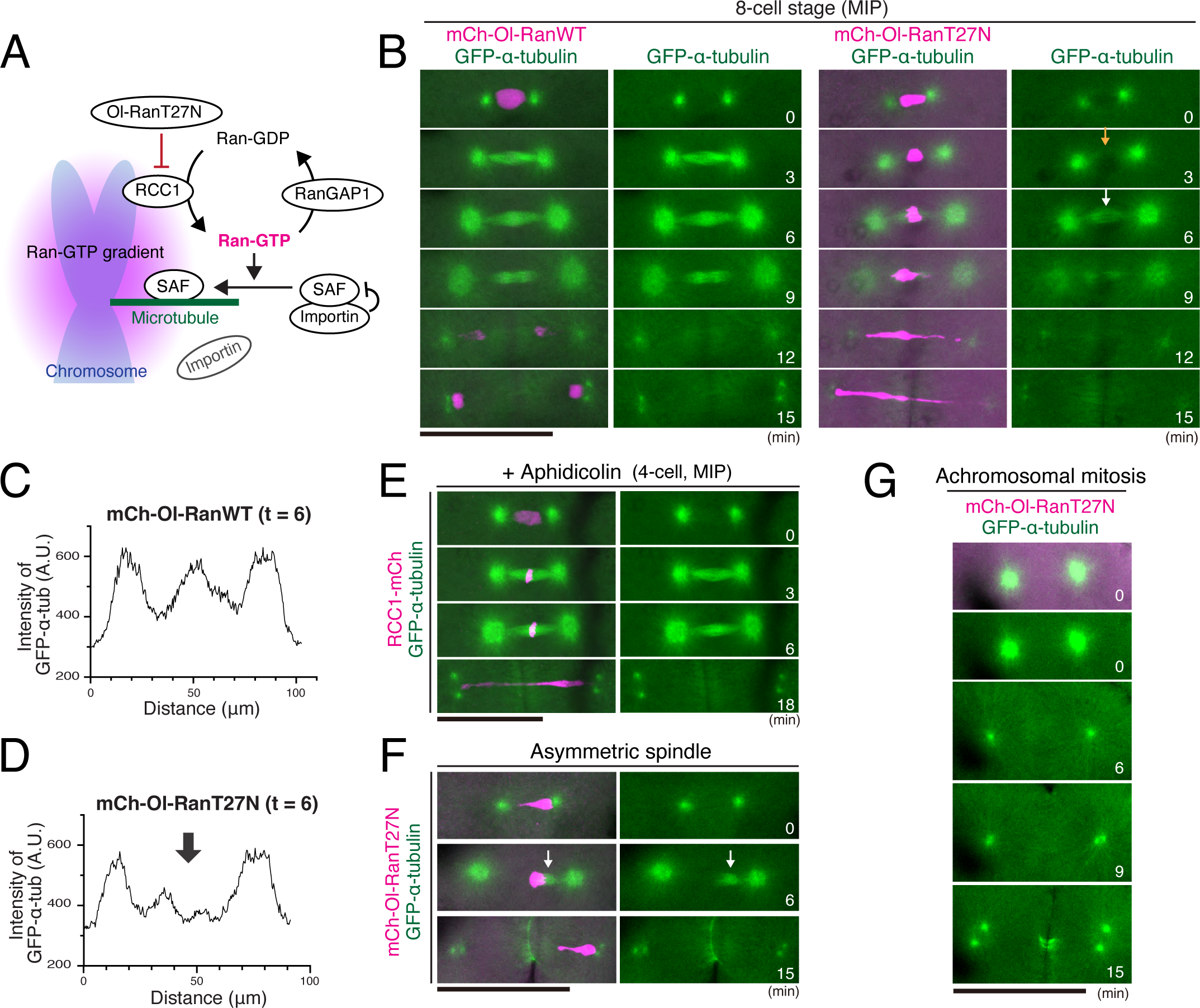
Expression of Ol-RanT27N causes spindle assembly defects followed by abnormal chromosome segregation. (A) Schematic representation of Ran-GTP-mediated spindle assembly by activated SAFs. (B) Live-cell images of 8-cell stage embryos showing normal spindle assembly in control (left) and abnormal spindle formation followed by severe chromosome mis-segregation in Ol-RanT27N expressing embryos (right). (C, D) Graphs of relative fluorescence intensities for line scans of Ol-RanWT (C) or Ol-RanT27N (D) expressing metaphase spindles shown in (B). An arrow in D indicates the decrease of MT intensity at the spindle midplane. (E) Live-cell images of 4-cell-stage spindles showing normal spindle assembly in aphidicolin-treated embryos despite abnormal chromosome segregation. (F, G) Live-cell images of Ol-RanT27N expressing embryos showing asymmetric spindle formation (F) and no spindle formation in achromosomal mitosis (G). Scale bars = 100 μm.

Ol-RanT27N expression also caused deformation or elongation of interphase nuclei (Figure 5B t=0, S5D t=-3), likely due to defects in nuclear envelop reformation and/or nucleocytoplasmic transport during interphase^57^. Previously, a similar abnormal chromosome segregation phenotype was observed^58^ by inhibiting DNA replication with aphidicolin, a DNA polymerase α inhibitor (Figure 5E t=18). However, aphidicolin-treatment did not affect spindle assembly timing or structure, or chromosome alignment (Figure 5E t=3, 6). Together, these data indicate that Ran-GTP promotes spindle MT nucleation during prometaphase and formation of a specialized MT network at the spindle midplane during metaphase in medaka embryos, although we cannot exclude the possibility that some interphase defects caused by Ol-RanT27N expression also contribute to its depletion phenotypes in mitosis.

### Centrosomes are sufficient for cleavage furrow formation, but not for spindle assembly

After expression of Ol-RanT27N, we unexpectedly found that some interphase blastomeres showed precocious detachment of a centrosome from the nucleus (Figure S5D t=-3), which resulted in asymmetric nucleation of spindle MTs near the centrosome in the subsequent mitosis (Figure 5F t=6). This asymmetric spindle finally resulted in mis-segregation of an entire chromosome (Figure 5F t=15). Regardless of these abnormalities, blastomeres continued cleavage divisions, likely due to the lack of cell cycle and mitotic checkpoints (Figure S5D and S5E). Interestingly, blastomeres without chromosomes (Figure 5F t=15) entered the next mitosis, in which the remaining two centrosomes were insufficient to assemble a spindle structure (Figure 5G t=0 and S5E t=6 and 30), but sufficient to form a cleavage furrow between them (Figure 5G t=15, and S5E t=24). These results indicate that chromosomes are required for spindle assembly in medaka early embryos, and that Ran-independent chromatin pathways collaborate with centrosomes in spindle MT nucleation near centrosomes (Figure 5F).

### AID2-mediated degradation of endogenous RCC1 disrupts functional spindle assembly and chromosome segregation in medaka early embryos

RanT24N interacts not only with RCC1, but also with importin-β, implying that RanT24N does not act solely as an RCC1 inhibitor^33^. To further validate our results, we next sought to disrupt Ran-GTP gradients by depleting RCC1 using auxin inducible degron 2 (AID2) technology^44^ (Figure 6A). As we did for mCherry knock-in, we integrated mAID-mClover-3×FLAG (mACF) coding sequence into the genome at the C-terminal region of the RCC1 gene (Figure S6A). Homozygous knock-in embryos grew normally.

**Figure 6.**
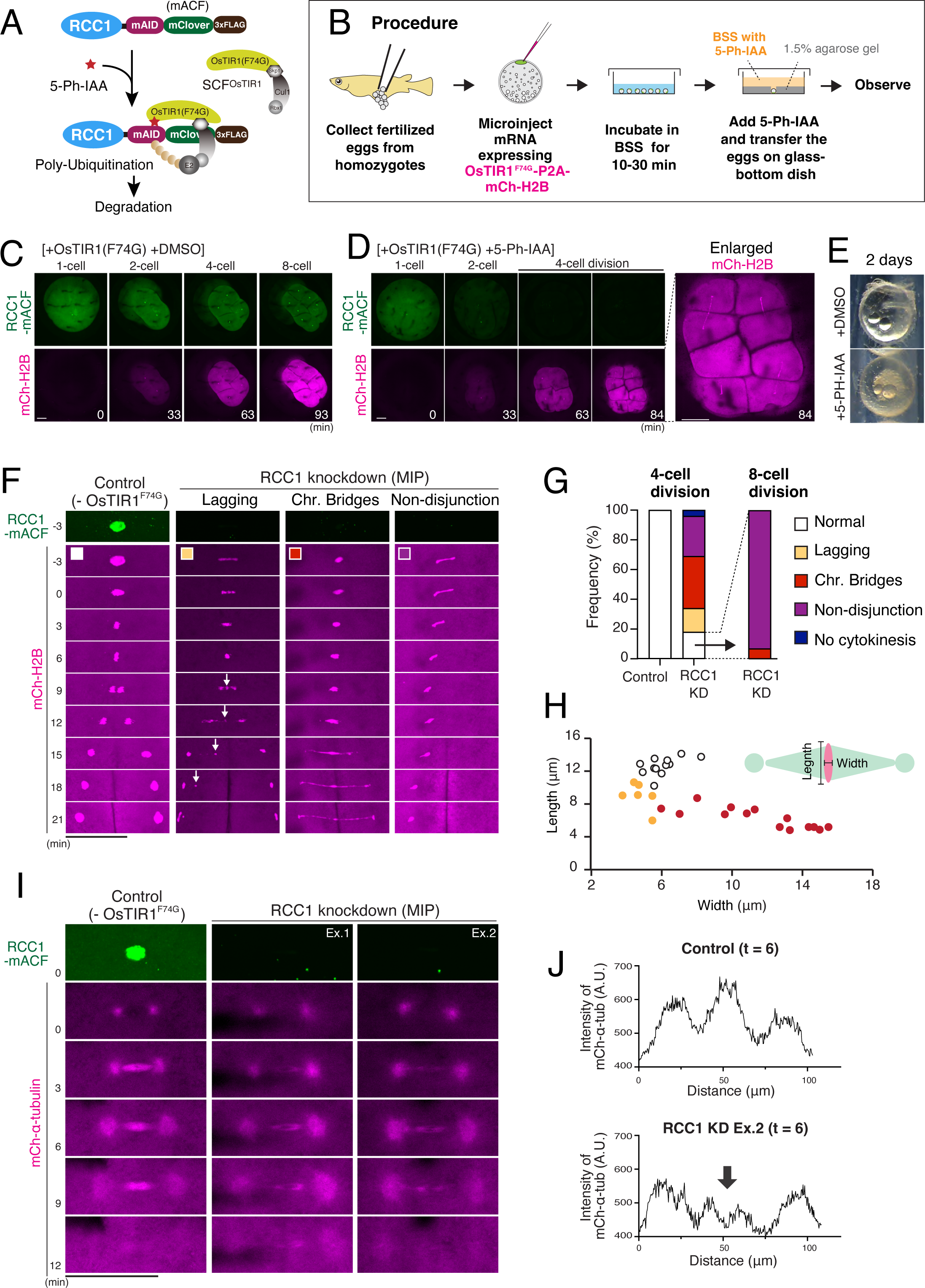
AID2-mediated degradation of endogenous RCC1 causes severe defects in spindle formation and chromosome segregation in medaka early embryos. (A) Schematic representation of AID2-mediated RCC1 degradation. (B) Procedure of AID-mediated protein knockdown in medaka fertilized eggs. (C, D) Representative live-cell images showing the fluorescence of RCC1-mACF and mCh-H2B in control (C) and 5-Ph-IAA treated (D) embryos. An enlarged image showing abnormal chromosome segregation is shown in D (right). (E) Phase contrast images 2 days after mRNA injection and treatment with DMSO (top) or 5-Ph-IAA (bottom) showing lethality in an RCC1-depleted embryo (bottom). (F) Live-cell images of control (left) and RCC1-depleted (3 columns on the right) 4-cell blastomeres showing chromosome segregation phenotypes: normal (white), lagging chromosomes (yellow), chromosome bridges (red), and chromosome nod-disjunction (purple). Anaphase lagging chromosomes (arrows) result in micronucleus formation. (G) Quantification of the frequency of abnormal chromosome segregation in control (n=28) and RCC1 knockdown 4-cell blastomeres (n=49) (left). Most daughter cells from normally dividing RCC1-depleted 4-cell blastomeres (n=14, white) show a non-disjunction phenotype in the subsequent 8-cell division (right). (H) Width and length of metaphase plates in G are plotted as colored circles for individual 4-cell spindles in control (white, n=12), and metaphase cells with lagging chromosomes (yellow, n=6) and chromosome bridging (red, n=14). (I) Live-cell images of control (left) and RCC1-depleted (2 columns on the right) 4-cell blastomeres showing disruption of the dense MT network at the spindle midplane in RCC1-depleted cells. (J) Graphs of relative fluorescence intensities for line scans of mCh-α-tubulin in (I), showing a decrease of mCh-α-tubulin intensity at the spindle midplane in RCC1-knockdown spindles. Scale bars = 100 μm.

To perform AID2-mediated protein knockdown, we collected naturally fertilized eggs from homozygous knock-in pairs and injected mRNAs encoding OsTIR1(F74G)-P2A-mCherry-Histone H2B (mCh-H2B) into one-cell stage embryos (Figure 6B, see Methods for details). mCh-H2B was used as an indicator of OsTIR1 expression as well as a marker of chromosomes. RCC1-mACF signals were visible in control embryos (Figure 6C), but decreased in response to expression of mCh-H2B in 5-Ph-IAA treated embryos, confirming efficient inducible degradation of RCC1 (Figure 6D). Although mCh-H2B expression levels were slightly variable among embryos, RCC1-mACF fluorescence reached an undetectable level before the third (4-cell) division in most embryos (Figure 6D). ∼80% (n=49) of 4-cell blastomeres depleted of RCC1 showed abnormal chromosome segregation phenotypes (Figure 6D t=84), and all embryos (n=14) showed embryonic lethality (Figure 6E). In the absence of OsTIR1(F74G), 5-Ph-IAA itself had no effects on RCC1 degradation and embryonic development (Figure S6B). In addition, when heterozygous embryos were used, embryos grew normally, even in the presence of OsTIR1(F74G) and 5Ph-IAA (Figure S6C and S6D), suggesting that half the normal amount of RCC1 is sufficient for embryonic development.

Homozygous RCC1-depleted blastomeres displayed lagging chromosomes, chromosome bridges, or chromosome non-disjunction during 4-cell divisions (Figure 6F and 6G). Phenotypic severity appeared to be correlated with the expression level of OsTIR1(F74G), indicated by the intensity of mCh-H2B (Figure 6F). ∼20% (n=49) of RCC1-knockdown (KD) blastomeres showed normal chromosome segregation at the 4-cell stage, likely due to residual RCC1, but all of these daughter cells displayed severe defects in the subsequent 8-cell division (Figure 6G). RCC1-knockdown blastomeres showed wider metaphase plates (Figure 6H), suggesting impaired chromosome-microtubule interaction (Figure 6H).

To analyze spindle structure in RCC1 knockdown embryos, we next expressed OsTIR1(F74G) and mCherry-α-tubulin. As observed in Ol-RanT27N-expressing embryos (Figure 5B), RCC1 knockdown caused slow spindle assembly during prometaphase (Figure 6I t=3). In addition, RCC1-depleted spindles failed to form the dense MT network around the spindle midplane (Figure 6I t=6, 6J, S6E), where spindle MTs surrounded chromosomes and built a hole-like structure (Figure S6E). RCC1 knockdown also caused asymmetric spindles (Figure S6F), as observed in Ol-RanT27N-expressing embryos (Figure 5F). Together, these results strongly support our finding that Ran-GTP gradients are critical to assemble functional spindles in medaka early embryos.

### RCC1 is dispensable for spindle assembly and chromosome segregation in blastula-stage embryos

In stark contrast to our results (Figure 6F), we previously reported that RCC1 depletion in HCT116 human cells does not result in abnormal chromosome segregation phenotypes^34^. To analyze the requirement of Ran-GTP in later-stage live embryos, we next added 5-Ph-IAA 8 hr post-OsTIR1(F74G) injection (hpi), which corresponds to blastula stage (Figure S2A). In the presence of both OsTIR1(F74G) and 5-Ph-IAA, RCC1-mACF signals were diminished and reached an undetectable level within 45 min (Figure 7A). To monitor chromosomes with mCh-α-tubulin, H2B fused with miRFP670-Nano3^59^ was used, which produces bright signals in later stages (Figure 7A-C), but not in early embryos (data not shown). In RCC1-knockdown blastomeres, bipolar spindles were still assembled (Figure 7B), but they were slightly shorter and appeared to have less intense microtubules around chromosomes (Figure 7B-7D) as observed in RCC1-depleted cultured human cells^34^. Mitosis was slightly prolonged (Figure 7E), but abnormal chromosome segregation was rarely observed (Figure 7F). Instead, RCC1-depleted blastomeres failed to form round nuclei with decondensed chromosomes after mitotic exit (Figure 7G), which most likely caused lethality of RCC1-depleted embryos. These data indicate that Ran-GTP becomes non-essential for functional spindle assembly and chromosome segregation, but remains indispensable for nuclear envelope reformation after mitotic exit in later-stage embryos.

**Figure 7.**
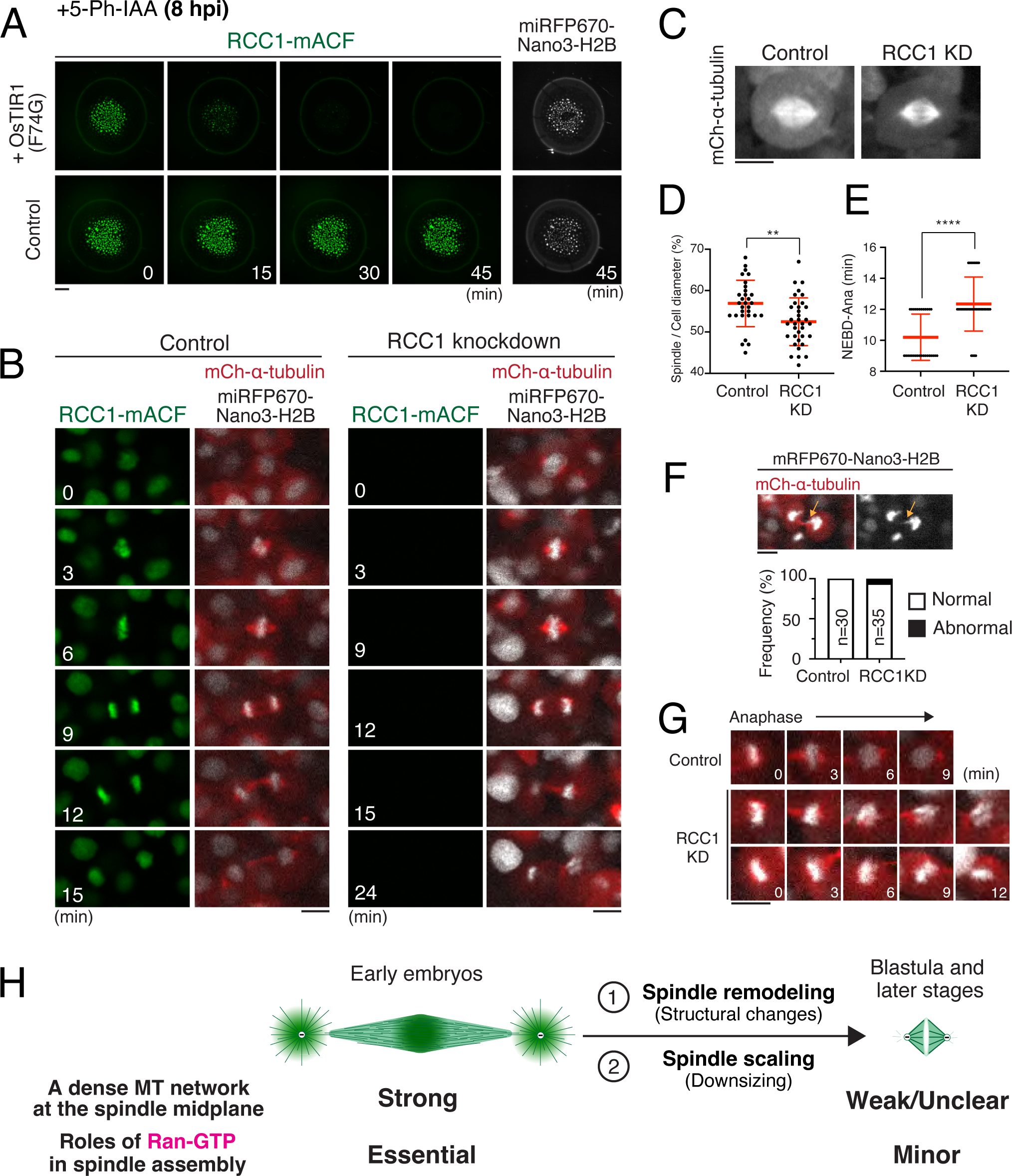
AID2-mediated RCC1 depletion in blastula embryos causes minor defects in spindle assembly and chromosome segregation. (A) Live-cell images showing the fluorescence of RCC1-mACF and miRFP6700-Nano3-H2B in control (lower) and OsTIR1(F74G)-expressing (upper) embryos. 5-Ph-IAA was added 8 hr post-injection (hpi). (B) Live-cell images showing the fluorescence of RCC1-mACF, mCh-α-tubulin, and miRFP6700-Nano3-H2B in control (left) and OsTIR1(F74G)-expressing (right) embryos. (C) Live images of mCh-α-tubulin in control and RCC1-depleted metaphase cells. (D) Ratio of spindle length and cell diameter in control (n=30) and RCC1 knockdown (n-35) cells. (E) Scatterplots of mitotic duration in control (10.2 ± 1.5, n=30) and RCC1-depleted (12.3 ± 1.7, n=35) cells. (F) Top: Live images of abnormal chromosome segregation (arrows). Bottom: Quantification of normal and abnormal anaphase in control and RCC1-depleted cells. (G) Representative live cell images showing nuclear reformation defects in RCC1 knockdown cells. (H) Diagram showing remodeling of spindle architecture and the requirement for Ran-GTP in early and later-stage embryos. See text for details. Error bars indicate mean ± SD; **p<0.01, **** p<0.0001. Scale bars = 100 μm (A) and 10 μm (B, C, F, G).

## Discussion

In this study, we established live functional assay systems in medaka early embryos. The efficiency of AID2-mediated protein degradation in medaka embryos (Figure 6D and 7A) is similar to that in cultured human cells^44^. Analyzing *in vivo* functions of other key conserved mitotic genes would further provide useful information about spindle assembly, positioning, and remodeling during early embryogenesis in medaka and other vertebrates.

In contrast to bovine and human zygotes^16, 17^, our live imaging revealed that early embryonic divisions in medaka are not error-prone despite very rapid mitoses (<12 min, Figure 2B and 4A) and a lack of a functional spindle assembly checkpoint (Figure 4F). Importantly, we found that medaka embryonic spindles have a specialized MT network at the spindle midplane in early stages (Figure 3A), which is assembled during metaphase (Figure 4A) in a Ran-dependent manner (Figure 5B and 6I) to achieve proper chromosome segregation (Figure 5B and 6F). Since centrosomes are largely separated from the spindle midplane (> 30 μm) after metaphase (Figure 4A), the chromosome-derived Ran-GTP gradient should act as an essential factor to assemble functional spindles for chromosome segregation in early embryos, like acentrosomal spindle assembly in female meiosis^31–33^. A dense midplane MT network is observed in zebrafish early embryonic spindles^7^, and *Xenopus laevis* stage 3 spindle intermediates^3^, but not clear in bovine 1- or 2-cell spindles^16, 60^. It is tempting to speculate that up-regulation of this mechanism may rescue the error-prone nature of bovine and human embryos, as recently shown for exogenous KIFC1 introduction to rescue a spindle instability phenotype in human oocytes^61^. It will be important to investigate how Ran-GTP assembles the midplane spindle structure and how it achieves faithful chromosome segregation in medaka and other vertebrate embryos.

On the other hand, Ran-GTP also contributes to the initial step of embryonic spindle assembly (Figure 5B t=3, and 6I t=3), in which Ran-GTP may promote MT nucleation by activating TPX2 and augmin dependently or independently of centrosome-dependent MT nucleation^62–65^. In addition, considering that weak spindle structures are finally formed in RCC1-deficient embryos (Figure 5B t=6, and 6I t=6), and that no spindles form in the absence of chromosomes (Figure 5G), other chromatin pathways such as the chromosome passenger complex^66–69^ may also function to assemble spindles in early embryos. It will also be interesting to analyze how Ran-independent pathways function with centrosomes in a distance-dependent manner (Figure 5F and S6F). These Ran-independent signals and centrosomes should be sufficient to form bipolar spindles in smaller cells after blastula stage (Figure 7B), likely owing to the reduced requirement for the midplane MT network and the short distance between chromosomes and centrosomes (Figure 7H).

In summary, this study provides the foundation to understand mechanisms of chromosome segregation and spindle assembly, positioning, and remodeling in medaka early embryos. Studying their detailed mechanisms is important to identify general and species-specific strategies that successfully coordinate chromosome segregation with cell differentiation and embryogenesis in vertebrates.

## Acknowledgments

We thank Kiyoshi Naruse, and the OIST animal resource section for technical assistance and support, and Marvin van Toorn for critical reading of the manuscript. We are grateful to NBRP Medaka (https://shigen.nig.ac.jp/medaka/) and NBRP Paramecium Laboratory (http://nbrpcms.nig.ac.jp/paramecium/) for providing OK-Cab (Strain ID: MT830) and *Paramecium* strain (PS000001A), respetively. This work was supported by grants from JSPS KAKENHI (17H05002 and 21H02481 to TK, and JP21H0419 and JP23H04925 to MTK), JST FOREST (JPMJFR224O to TK) and CREST (JPMJCR21E6 to MTK), the Takeda Foundation (to TK), the Uehara Foundation (to TK), and the Okinawa Institute of Science and Technology Graduate University, Japan.

## Author contributions

Conceptualization, TK; Investigation, AK and TK; Formal analysis, AK and TK; Methodology, AK, TN, SA, MTK, MT, and TK; Rearing and maintenance of Medaka fish, AK, SH, and TK; Writing, TK; Supervision, TK; Funding Acquisition, TK. and MTK

## Methods

### - Fish maintenance

Fish experiments were conducted in accordance with protocols (2019-273-5, 2022-369) approved by the Animal Care and Use Committee at Okinawa Institute of Science and Technology Graduate University (OIST). The OK-Cab strain (MT830) of medaka (*Oryzyas latipes*) was obtained from the National Bio-Resource Project Medaka (NBRP Medaka) and used as the parental strain. Medaka were maintained in fresh water at 26-28°C under a regulated photoperiod (14-h light and 10-h dark). Medaka embryos were cultured at 27-28°C. Medaka larvae were grown in a tank without water circulation for 2-3 weeks and then transferred into a tank with water circulation (Meito system, Meito Suien or ZebTEC, Tecniplast). Fish were fed three times per day with live brine shrimp at 1 pm, and a commercial dry feed (Hikari-Lab, Kyorin) around 9 am and 5 pm. For larvae, paramecia (PS000001A), provided by NBRP Paramecium Laboratory, were provided with dry feed for the first 3-5 days. From 4-6 days, live brine shrimp were fed instead of paramecia. The particle size of dry feed (hikari-lab, 130,270,450) was adjusted according to fish size.

Naturally fertilized healthy eggs were used for imaging. Fluorescence of RCC1-mCh and RCC1-mACF was constantly observed in all heterozygous and homozygous knock-in embryos. However, fluorescence of EGFP-α-tubulin was variable, even in homozygous embryos and was sometimes undetectable for unknown reasons. We selectively used embryos having similar fluorescence intensity of EGFP-α-tubulin in all experiments. Established plasmids, Medaka strains, and sequence information about guide RNA (gRNA) and PCR primers used in this study are described in Table S1,S2,S3 and S4, respectively.

### - Plasmid Construction

Donor dsDNAs for CRISPR/Cas9-mediated genome editing (Figure S1A and S6A) were constructed according to the protocol of Gutierrez-Triana et al.,^45^. To design a donor plasmid, PAM sequences were searched around the stop codon of the medaka (*Oryzias latipes*) RCC1 gene (Ol-RCC1) on chromosome 16 using CCTop, CRISPR/Cas9 target online predictor (https://cctop.cos.uni-heidelberg.de:8043/)^70^. Silent mutations were introduced to the gRNA target and PAM sequences. The stop codon was mutated to code glycine (G), and a BamHI site was introduced in-frame after the G so that CDS cassettes flanked by BamHI could be inserted. The designed DNA containing these mutations, a BamHI site and homology arms for Ol-RCC1 (∼450-bp homology arms) was synthesized by a gene synthesis service (Genewiz, South Plainsfield, NJ). mCherry2 and mAID-mClover-3×FLAG(mACF) cassettes were created using pMK281 (addgene #72797) and pTK398 (addgene #114714), respectively, and inserted at the BamHI site on the synthesized DNA plasmid to make donor plasmids, pTK997 and pTK1023. Using these plasmids as a template, dsDNAs were amplified by PCR using modified primers with 5’Bioton – 5 x phosphorothioate bonds (synthesized by eurofins) and PrimeSTAR Max (Takara).

To construct a knock-in (KI) donor vector for EGFP-α-tubulin (Figure S1B), Mbait-hsp-EGFP-BGHpA plasmids^71^ were linearized by PCR using KOD-Plus-Neo (TOYOBO), followed by insertion of a flexible linker^72^ at the C-terminus of EGFP using the InFusion system (Takara). α-tubulin CDS was amplified from a cDNA clone, olli34h17, (gifted by NBRP medaka), and inserted into the XbaI site of the resultant Mbait-hsp-EGFP-FL-BGHpA plasmid to generate Mbait-hsp-EGFP-FL-α-tubulin-BGHpA (KI vector). A CRISPR/Cas9 target site at 5’UTR of α-tubulin was searched using CCTop.

To exogenously express EGFP- or mCherry-fusion proteins, CDS of EGFP-EB1 (addgene #46364) and EMTB-3×GFP (addgene #26741) were inserted between BamHI-SnaBI sites in pCS2+hSpCas9 vector (addgene #51815, a gift from Dr. Kinoshita in Kyoto University). H2B CDS was obtained from pSNAPf-H2B (New England Biolabs). CDS of Ol-RanT27N and miRFP670-Nano3 was synthesized by eurofin, and Ol-RanWT CDS was created by PCR-based mutagenesis.

To make OsTIR1-P2A-mCh-H2B and OsTIR1-P2A-mCh-α-tubulin plasmids, pMK411 (addgene #140659) was used as a template to amplify the CDS of OsTIR1-P2A, which was fused with mCh-H2B or mCh-α-tubulin by PCR. CDSs were inserted between BamHI-SnaBI sites of the pCS2+hSpCas9 vector.

### - gRNA synthesis

For synthesis of gRNA, T7 tagged DNA templates containing the gRNA sequence were amplified by PCR, as described previously^73^. For synthesis of Mbait gRNA, a DR274 vector (addgene # 42250) containing Mbait (gifted by Dr. Yasuhiro Yamamoto at Osaka Medical and Pharmaceutical University) was used as a template to amplify fragments containing T7 and gRNA sequences by PCR followed by gel purification. PCR fragments were used as a template for *in vitro* transcription with a MEGAscript T7 kit (Thermo Fisher Scientific, AM1333) according to the manufacturer’s instructions. Synthesized gRNAs were purified by ammonium acetate precipitation.

### *In vitro* transcription of mRNA

For synthesis of Cas9 mRNA, the pCS2+hSpCas9 vector was linearized by NotI digestion, followed by *in vitro* transcription using the mMessage mMachine SP6 kit (Thermo Fisher Scientific, AM1340) according to the manufacturer’s instructions. The synthesized RNA was purified with an RNeasy Mini kit (Qiagen). To synthesize other mRNAs, template plasmids were linearized with NotI or BssHII.

### - Microinjection

To generate RCC1-mCh (Figure S1A) or RCC1-mACF (Figure S6A) knock-in strains, 50 ng/µL gRNA targeting RCC1, 150 ng/µL Cas9 mRNA and 10 ng/µL dsDNA were injected into one-cell stage medaka embryos. To establish an EGFP-α-tubulin knock-in strain (Figure S1B), 25 ng/µL Mbait gRNA, 25 ng/µL gRNA targeting α-tubulin, 50 ng/µL Cas9 mRNA and 2.5 ng/µL KI vector were used. To exogenously express EGFP- or mCherry-fusion proteins, 150 ng/µL mRNAs were injected into one-cell embryos.

Glass needles were made from borosilicate glass capillaries (Model No: G100F-4, Order No: 64-0787, Warner Instruments) using a needle puller (PC-100, Narishige), and attached to a capillary holder connected with a microinjector Femto Jet 4i (eppendorf). Needles were manually controlled by a micro-manipulator (MN-153, Narishige) on the stage of a stereomicroscope (Leica M80).

### - Genotyping and selection of knock-in strains

Injected eggs were incubated at 27-28°C. Hatched G0 fish were grown until they exceeded 1 cm in length. Small pieces of tail fin were cut and added to 20 µL lysis buffer (0.1M Tris-HCl, 0.2 M NaCl, 0.1% SDS, 5 mM EDTA). After 5 min incubation at 95°C, 80 µL H_2_O were added. After centrifugation, 1-µL aliquots of supernatant were used for genomic PCR with KOD FX neo (Toyobo) and appropriate primers (Table S4). Candidate G0 fish were crossed with wild-type counterparts, and fertilized eggs were analyzed for fluorescence of RCC1-mCh or RCC1-mACF, 1 or 2 days after fertilization. Genomic DNA was extracted from some fluorescence-positive F1 embryos, and genomic PCR was performed. Proper knock-in of the construct was confirmed by direct sequencing of PCR products. Other positive embryos from the same G0 fish were reared, and F1 male and female fish were mated to obtain a homozygous F2 generation. PCR products were analyzed by normal gel electrophoresis or using an automatic microchip electrophoresis system (MCE-202 MultiNA: Shimazu, Kyoto, Japan).

To select EGFP-α-tubulin knock-in strain (Figure S1B), embryos with EGFP fluorescence in their eyes and throughout their bodies were raised as G0 founders and were crossed with wild-type medaka to obtain a stable F1 transgenic line. Insertion of the KI vector at the target site was examined by PCR using specific primers for the KI vector and the promoter of α-tubulin (upstream of gRNA target site).

### - Microscope System and live imaging

Imaging was performed by spinning-disc confocal microscopy with a 20× 0.95 numerical aperture objective lens (APO LWD 20X WI λS, Nikon, Tokyo, Japan). A CSU-W1 confocal unit (Yokogawa Electric Corporation, Tokyo, Japan) with three lasers (488, 561, 640 nm, Coherent, Santa Clara, CA) and an ORCA-Fusion digital CMOS camera (Hamamatsu Photonics, Hamamatsu City, Japan) were attached to an ECLIPSE Ti2-E inverted microscope (Nikon) with a perfect focus and a water-supply system. Images were captured using NIS-Elements software (Nikon). Samples were imaged at room temperature (24-25°C).

To hold living embryos for time-lapse imaging, a handmade device was created: 1.5 % agarose solution was added to glass-bottomed dishes (CELLview™, #627860, Greiner Bio-One, Kremsmünster, Austria), and a 7-unified cover-glass (18 mm x 18 mm with 0.12-0.17 mm thickness, Matsunami) was set as a mold in the center of the glass-bottomed dish. When the agarose became solid, the unified glass was removed to make a concave pocket in the agarose gel, 0.8-1.2 mm in width and ∼3 mm in height on the bottom glass (Figure 6B). 2-8 embryos were put in the pocket in a row with the blastodisc facing the glass (Figure 1A). The dish was filled with ∼2 mL medaka balanced salt solution (BSS, 0.65% NaCl, 0.04% KCl, 0.02% MgSO_4_・7H_2_O, 0.02% CaCl_2_・2H_2_O, sterilized and adjusted to pH 8.3 with 5% NaHCO_3_) without phenol red. 11-13 z-section images 5-μm in thickness were acquired every 3 min or 1 min with camera binning 1. Green and red fluorescence and DIC images were captured in this order with exposure times of 500 msec, 1 sec, 100 msec, respectively. X-Y-Z positions of 2-7 embryos were memorized and automatically imaged for 6-10 h in a time-lapse experiment. For figures, 8-bit maximally projected z-stack images (MIP) or single z-section images are shown as indicated. MIP images were created using NIS-Elemetns or Fiji. Signals were linearly adjusted using Fiji and Photoshop to optimize image clarity, and images were arranged using Adobe Illustrator. Phase-contrast images of embryos for Figure 6E and S6D were taken with a Leica M80 stereo microscope and a Leica MC190 HD camera.

### - AID2-mediated protein knockdown

For AID2-mediated protein degradation, mRNAs encoding OsTIR1(F74G)-P2A-mCherry-H2B or OsTIR1(F74G)-P2A-α-tubulin were injected into one-cell embryos. Injected embryos were cultured in BSS without phenol red for 10-30 min with gentle shaking. The BSS was replaced with 2 mL BSS containing 10 μM 5-Ph-IAA or 0.1% DMSO, and then these embryos were transferred with the solution to the handmade glass-bottom dish. After adjusting the orientation of eggs in the agarose pocket, eggs were observed by microscope (Figure 6B). For experiments in Figure 7, injected embryos were cultured in medaka BSS for 8hr before 5-Ph-IAA treatment.

### - Quantification of Fluorescent intensities and phenotypes

For measurement, metaphase spindles having both centrosomes/poles in a single-focal plane were selected. To quantify fluorescence intensities of EGFP-α-tubulin, EGFP-EB1, EMTB-3×GFP, and mCh-α-tubulin, line scans were performed in Fiji. A line 5-pixels wide was drawn on single-z-section 16-bit images (Figure 3B, 3F, 3G, 4C) or MIP images (Figure 5C, 5D, 6J) as it passed through both spindle poles and centrosomes (Figure 3B). Spindle length and width were measured using NIS-Elements software. Graphs were created using Prism 9 (GraphPad Software, La Jolla, CA) or Excel (Microsoft).

### - Statistical Analysis

Mean, SD, and statistical significance using two-sided Welch’s t-tests were determined using GraphPad Prism version 9 (GraphPad Software, La Jolla, CA). P values are shown as *: **: p<0.01 and ****: p<0.0001.

## Supplementary Information

### Supplementary Figure Legends

**Figure S1.**
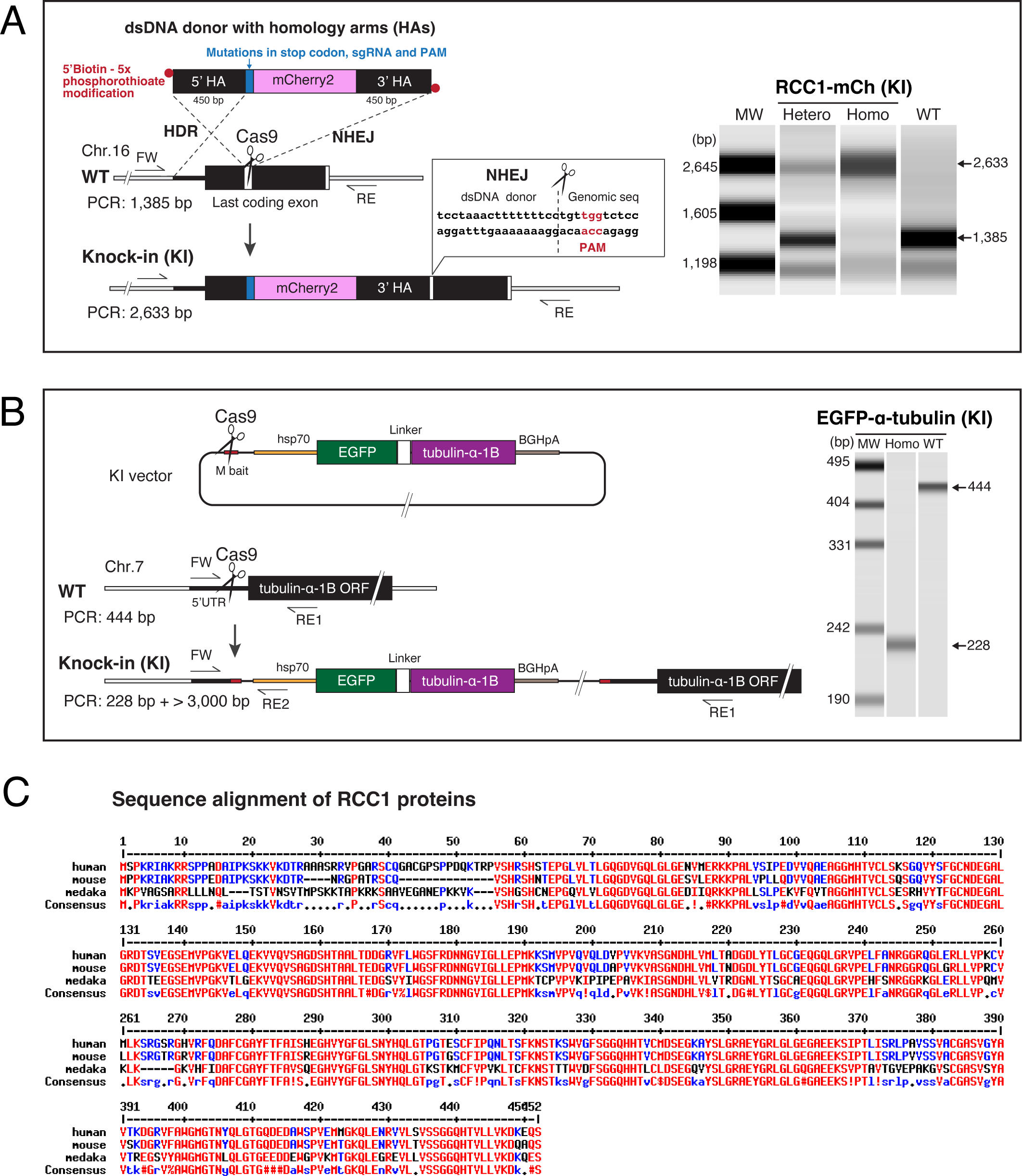
Generation of transgenic medaka strains. Related to Figure 1. (A) Left: Schematic representation of generation of the RCC1-mCherry (RCC1-mCh) knock-in (KI) strain using dsDNA as a donor. Although HDR-mediated knock-in was expected, the genomic PCR result of the candidate strain showed a longer band (∼2.6 kb). Sequence analyses of amplified fragments confirmed that the 3’ homology arm (HA) of the dsDNA donor was integrated into the genome via NHEJ, but not HDR. Right: PCR-based genotyping of the *RCC1* gene in the parental wild-type (WT) and KI strains. A single band of around 2.6 kb confirms homozygous insertion in the KI strain. (B) Left: Schematic representation of generation of the EGFP-α-tubulin KI strain. See Methods for details. Right: PCR-based genotyping of the EGFP-α-tubulin KI strain using three primers, FW, RE1 and RE2. A 228 bp band without a 444-bp band confirms homozygous insertion of the EGFP-α-tubulin construct. (C) Amino acid sequence alignment of RCC1 proteins in *H. sapiens* (NP_001041659), *M. musculus* (NP_001184011), and *O. latipes* (XP_023820025) using MultAlin (http://multalin.toulouse.inra.fr/multalin/)^1^.

**Figure S2.**
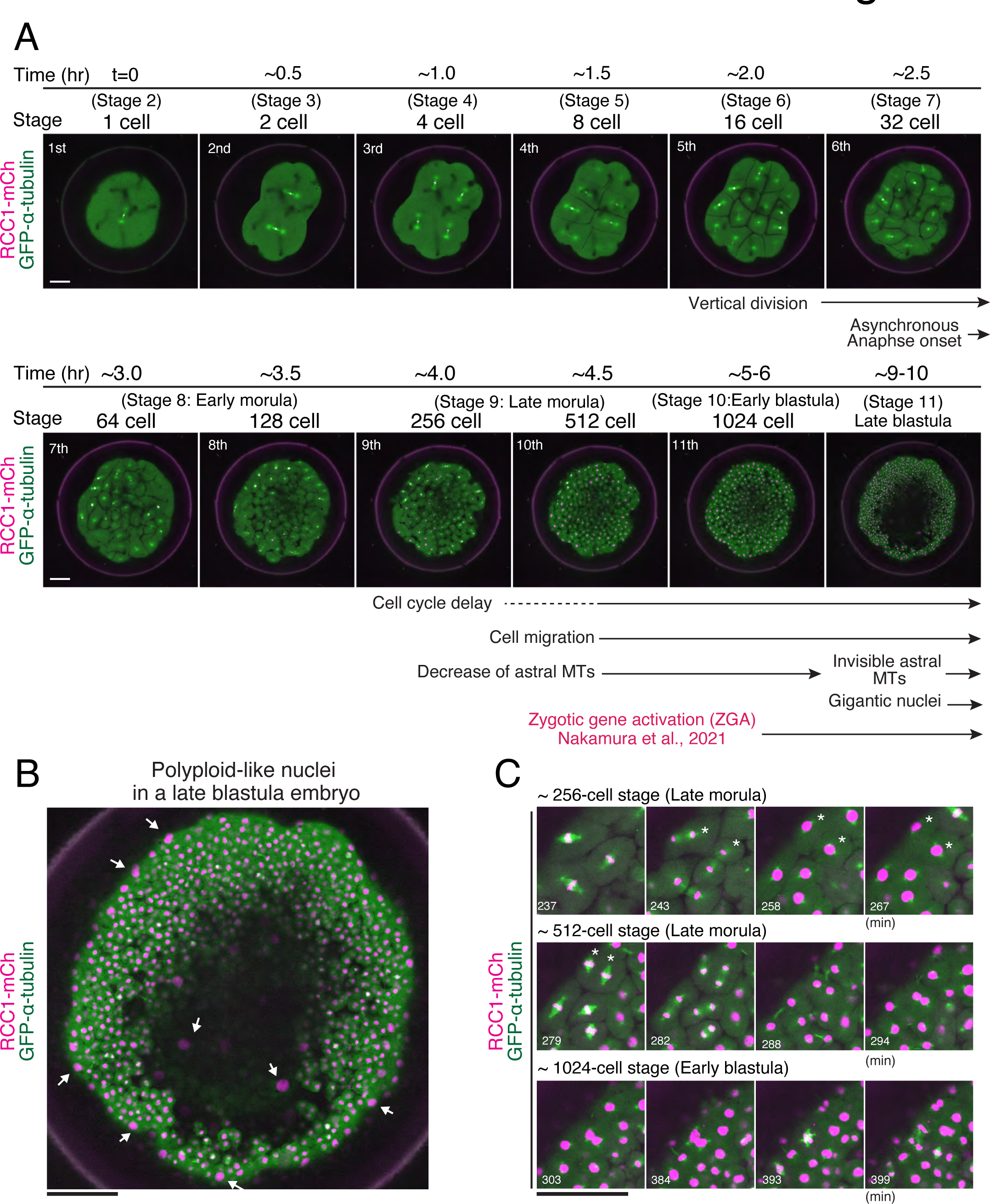
Live-cell images of early medaka embryos. Related to Figure 2. (A) Representative live-cell images showing metaphase spindles in indicated embryonic stages. Single z-section images are shown. Vertical divisions start from the 5^th^ division at the 16-cell stage. Structural changes of spindles roughly coincide with cell migration, cell cycle remodeling, and zygotic gene activation around early blastula stage. (B) A live image of a late blastula embryo showing a polyploid-like large nuclei formation (arrows) during normal development. (C) MIP images of the 3-z-sections showing two separated daughter cells (asterisks) were fused, resulting in generation of polyploid-like cells. Scale bars = 100 μm.

**Figure S3.**
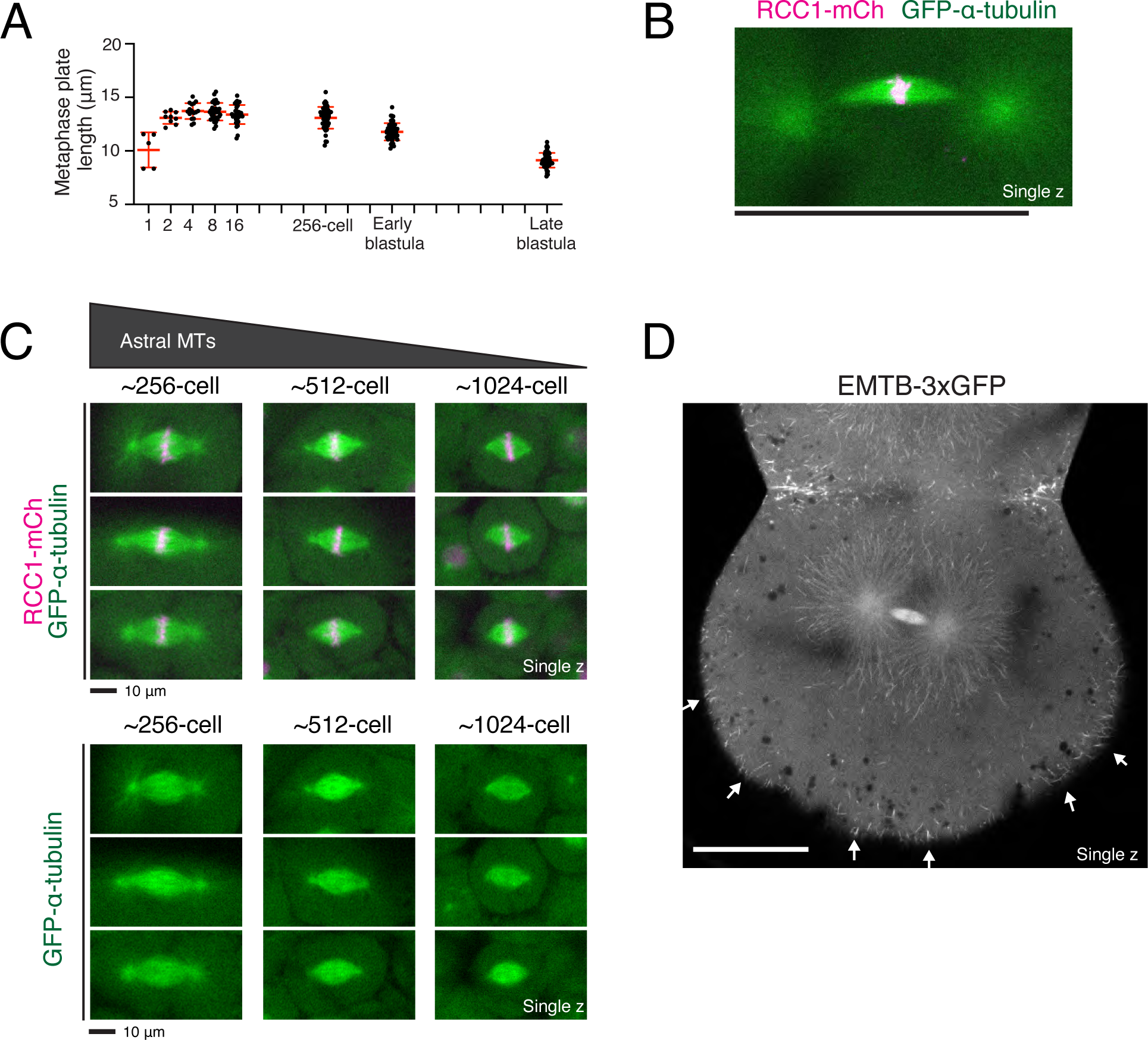
Morphological dynamics of medaka early embryonic spindles. Related to Figure 3. (A) Quantification of metaphase plate length in indicated stages. N= 5, >9, >15, >24, >24, >50, >50, >50 for 1-, 2-, 4-, 8-, 16-, 256-, early blastula, and late blastula stages, respectively, from 5 embryos. (B) A live fluorescent image showing a bent zygotic spindle. (C) Live metaphase-spindle images in ∼256-, ∼512-, and ∼1024-cell stage embryos showing a gradual decrease of astral microtubules. Single z-section images are shown. Scale bars = 10 μm. (D) A single z-section image of EMTB-3×GFP showing clear astral microtubules in a 2-cell embryo. EMTP-3×GFP also displayed microtubule-like signals around cortical cell membrane (arrows). Scale bars = 100 μm except for (C).

**Figure S4.**
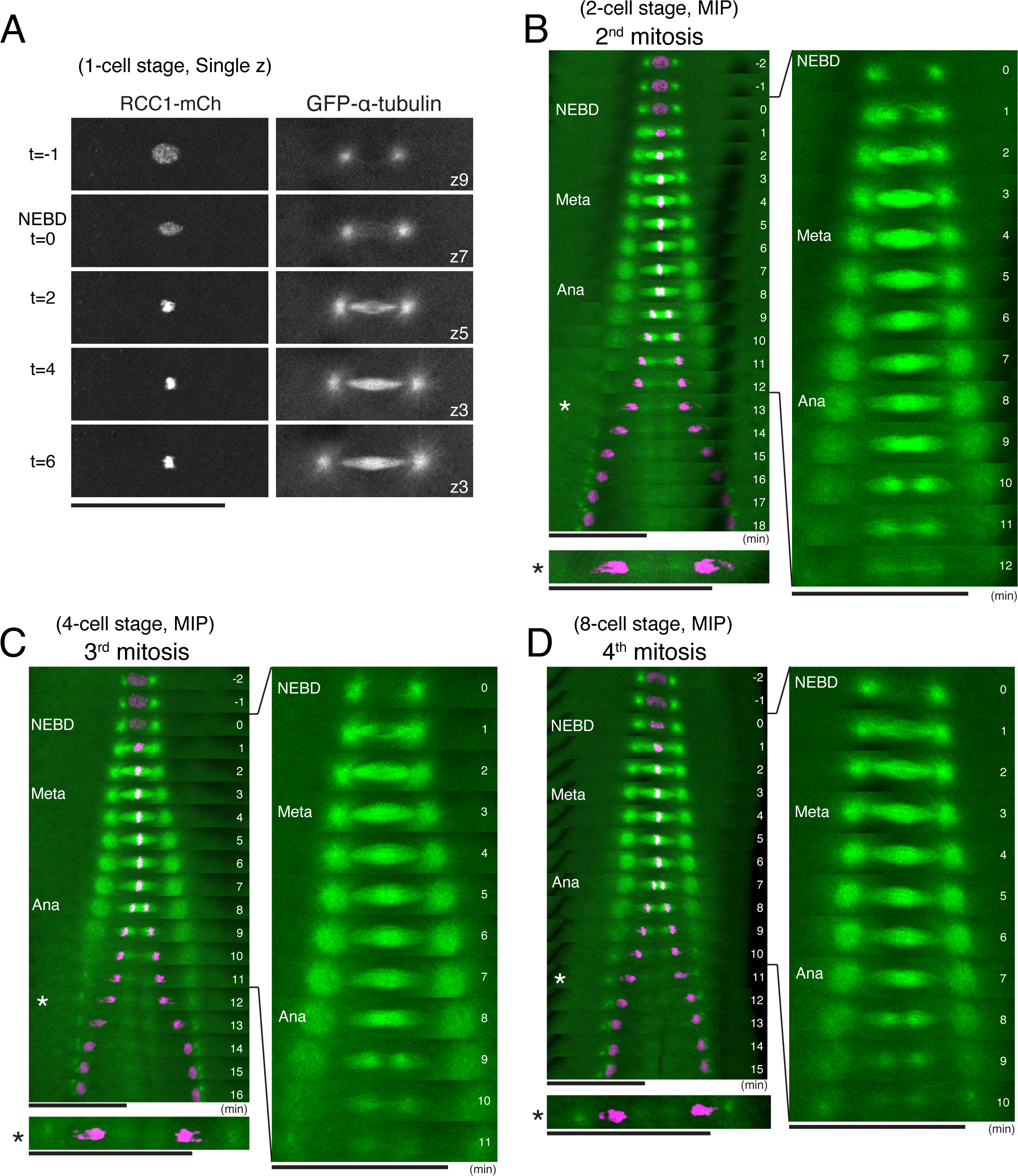
Assembly dynamics of a dense midplane MT network in early embryonic spindles. Related to Figure 4. (A) Live-cell, single z-section images during zygotic spindle assembly. Focal z-section positions change during embryonic spindle assembly. (B-D) Kymographs showing the second (B), third (C), and fourth (D) mitotic spindle assembly processes in a medaka embryo. A dense midplane MT network is formed during metaphase, as observed in the first mitosis (Figure 4A). Asterisks indicate a telophase chromosome or deformed nucleus migrating to centrosomes. Scale bars = 100 μm.

**Figure S5.**
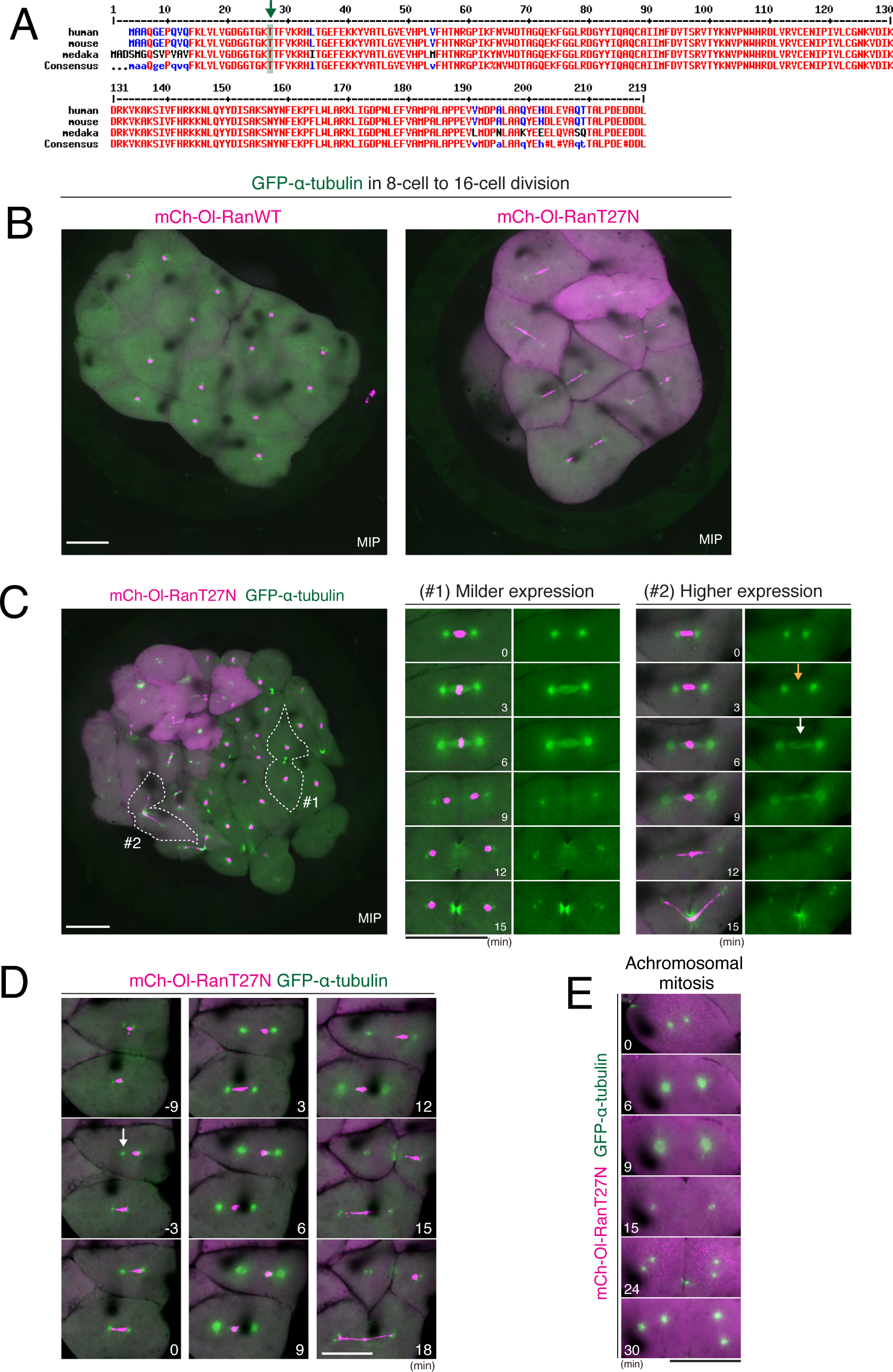
Mitotic phenotypes in RanT27N-expressing embryos. Related to Figure 5. (A) Amino acid sequence alignment of Ran proteins in *H. Sapiens* (NP_006316.1), *M. Musculus* (NP_033417.1), and *O. latipes* (XP_004073279.1) using MultAlin^1^. (B) Live-cell images of 8-cell-stage embryos showing normal chromosome segregation in control (left) and abnormal chromosome segregation in Ol-RanT27N-expressing embryos (right). (C) Left: a whole embryo image showing that mCh-Ol-RanT27 causes abnormal chromosome segregation in an expression-level-dependent manner. Right: time-lapse image sequences of cells showing milder (#1) or higher (#2) expression of mCh-Ol-RanT27N. A delay of spindle formation (a yellow arrow) and a defect in specialized midplane MT network formation were observed in (#2), but not (#1), cell. (D) Time-lapse image sequences showing that one of centrosomes detaches from the nucleus (t=-3, arrow), resulting in asymmetric spindle formation followed by asymmetric segregation of all chromosomes into one daughter cell. This generated a daughter cell lacking a nucleus. (E) Live images of achromosomal mitosis showing cytokinesis without assembling the spindle between centrosomes in medaka embryos. Scale bars = 100 μm.

**Figure S6.**
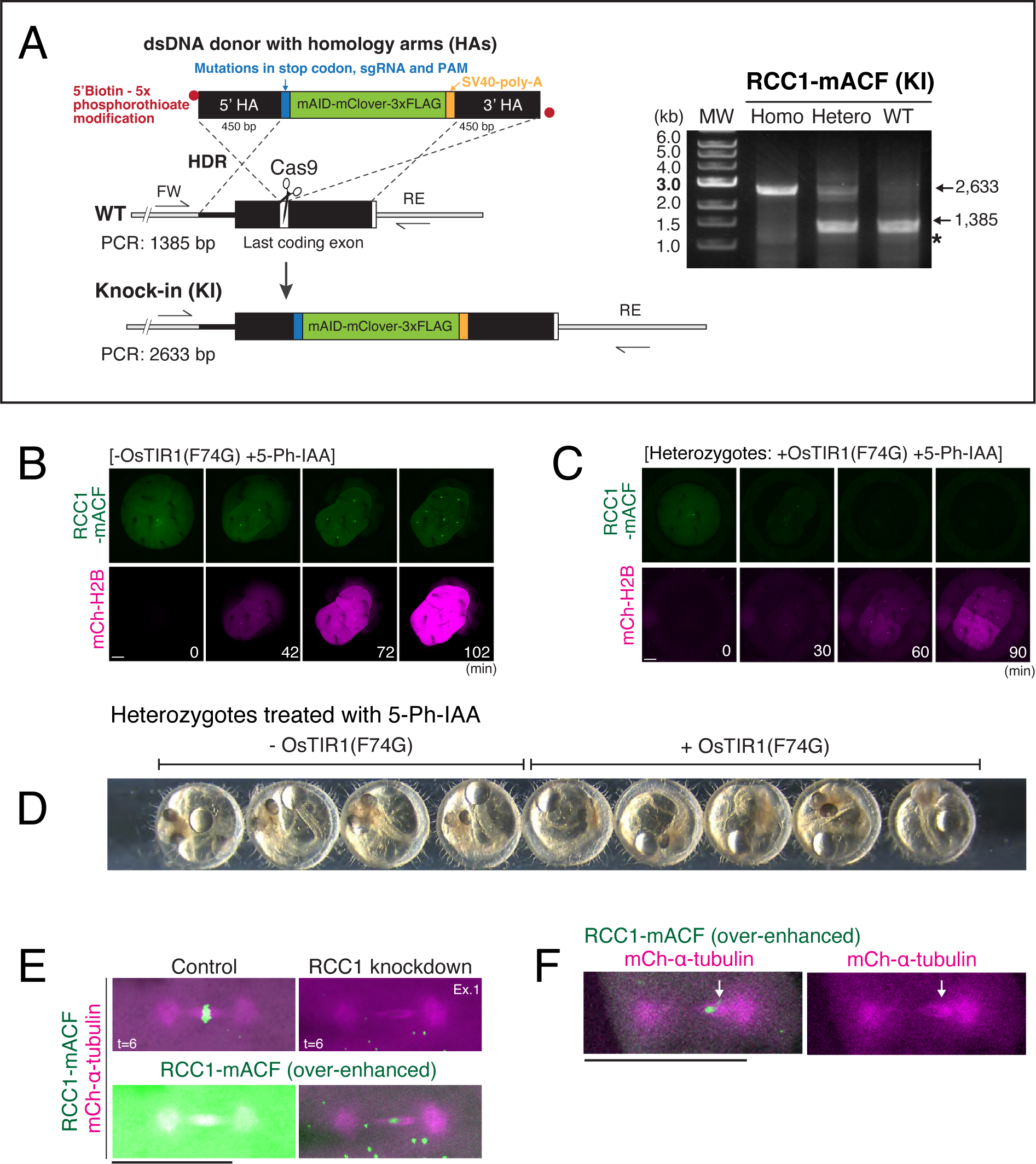
AID2-mediated depletion of RCC1 in medaka early embryos. Related to Figure 6. (A) Schematic representation of the generation of RCC1-mAID-mClover-3×FLAG (RCC1-mACF) knock-in (KI) strain using dsDNA as a donor. An SV40-polyA sequence is also integrated. Right: PCR-based genotyping of the *RCC1* gene in the parental wild-type (WT), and homozygous and heterozygous KI strains. A single band of around 2.6 kb confirms homozygous insertion in the KI strain. An asterisk indicates a non-specific band. (B, C) Representative live-cell images showing the fluorescence of RCC1-mACF and mCh-H2B in -OsTIR1(F74G) control (B), and heterozygous 5-Ph-IAA treated embryos expressing OsTIR1(F74G) (C). (D) Phase-contrast images of heterozygous RCC1-mACF embryos 4 days after mRNA injection and treatment with 5-Ph-IAA showing no developmental defects, regardless of the presence or absence of OsTIR1(F74G). (E) Top: live-cell images of control (left) and RCC1 protein-knockdown (right) 4-cell blastomeres showing disruption of the dense MT network at the spindle midplane in RCC1-depleted cells. Bottom: images with over-enhanced green fluorescence intensities showing remaining RCC1-mACF signals are located in the MT-less region in the RCC1-knockdown spindle. (F) Live-cell images of RCC1-knockdown embryos showing asymmetric spindle formation. Scale bars = 100 μm.

## Supplementary Movie

**Movie S1.** Related to Figure 2A. Time-lapse movie of MIP images of a fertilized medaka embryo showing chromosomes (RCC1-mCh) and microtubules (GFP-α-tubulin) during early embryonic divisions.

**Table S1:**
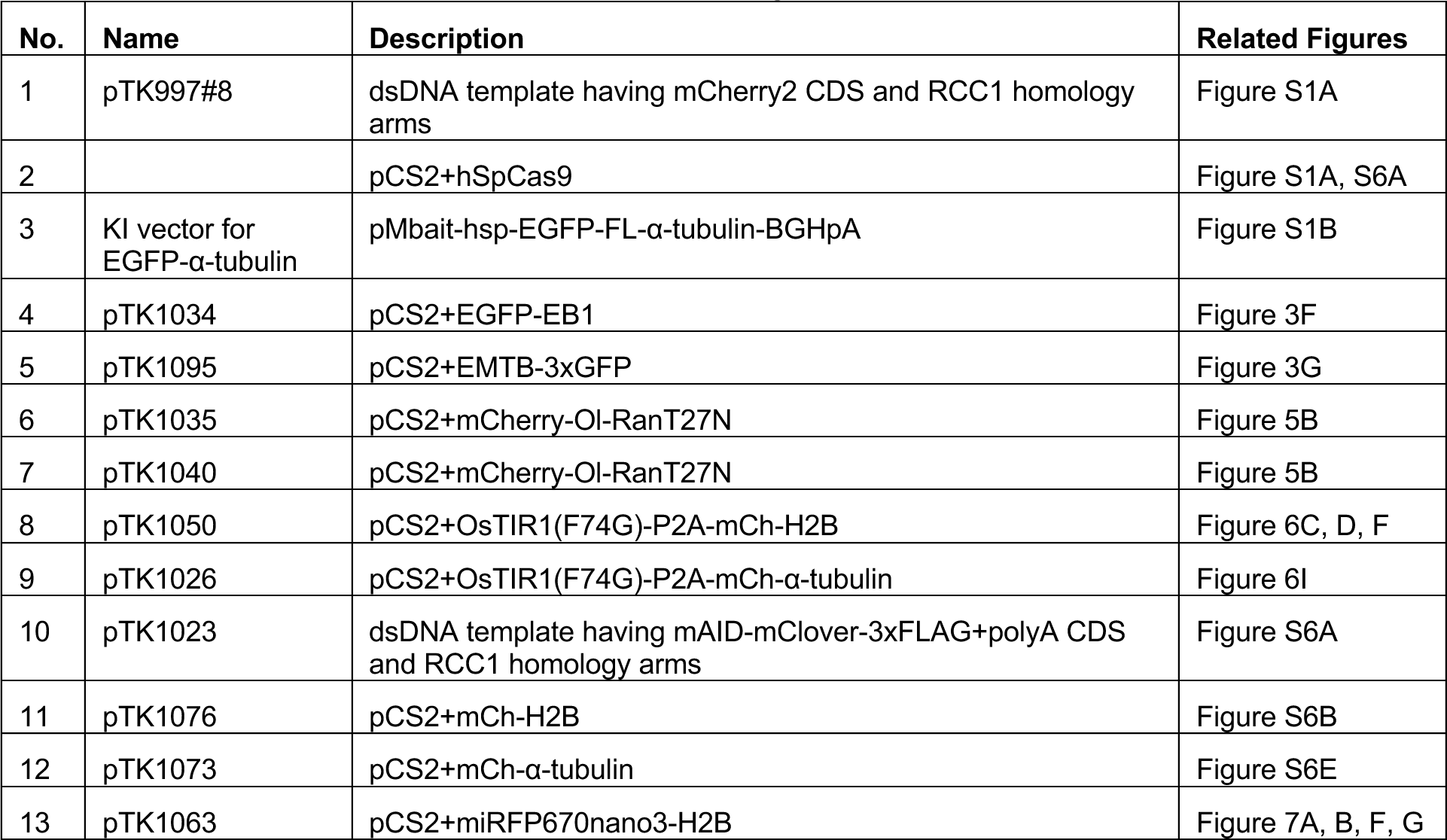
Plasmids established in this study.

**Table S2:** Medaka strains used in this study.

| No. | Name | Description | Plasmids used |
| --- | --- | --- | --- |
| 1 | OK-Cab (MT830) | National Bio-Resource Project Medaka |  |
| 2 | RCC1-mCh | A knock-in strain having mCherry at the RCC1 locus. | pTK997#8, pCS2+hSpCas9 |
| 3 | EGFP-α-tubulin | A knock-in strain having EGFP-α-tubulin at the tubulin-α-1B locus | KI vector for EGFP-α-tubulin, pCS2+hSpCas9 |
| 4 | RCC1-mCh + EGFP-α-tubulin | A double knock-in strain obtained by crossing No.2 with No.3 strains. | |
| 5 | RCC1-mACF | A knock-in strain having mAID-mClover-3xFLAG (mACF) at the RCC1 locus. | pTK1023, pCS2+hSpCas9 |

**Table S3:**
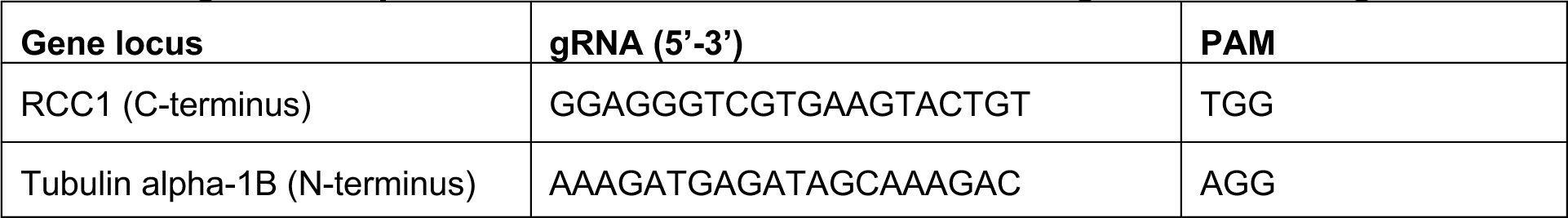
gRNA sequences for CRISPR/Cas9-mediated genome editing.

**Table S4:**
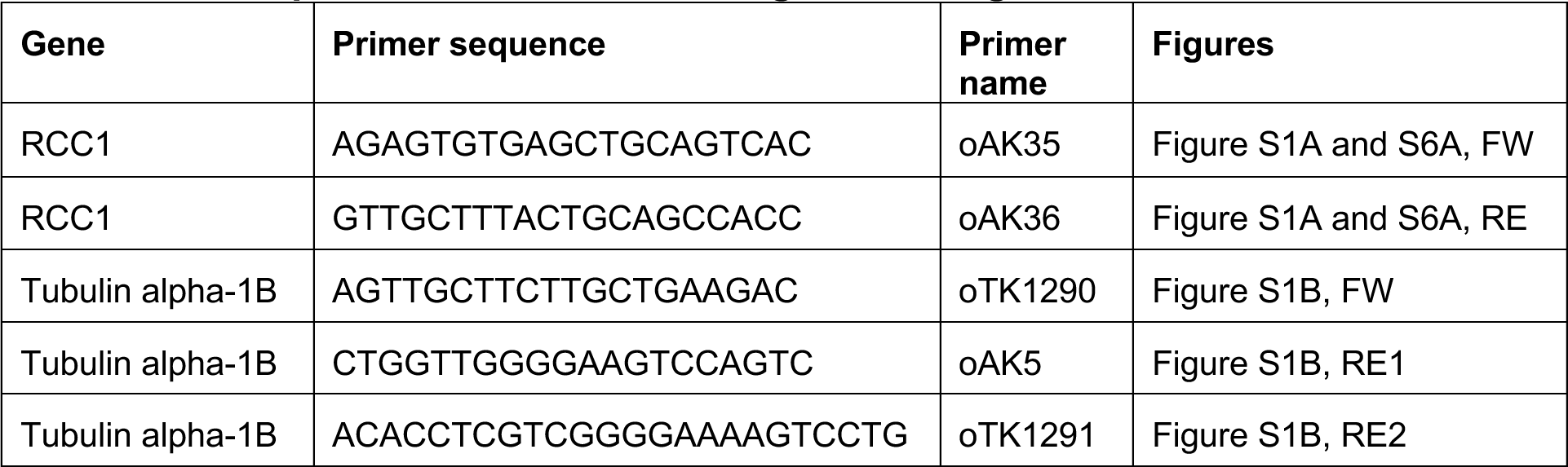
PCR primers used to confirm gene editing.

